# *C2cd6*-encoded CatSperτ Targets Sperm Calcium Channel to Ca^2+^ Signaling Domains in the Flagellar Membrane

**DOI:** 10.1101/2021.08.16.456347

**Authors:** Jae Yeon Hwang, Huafeng Wang, Yonggang Lu, Masahito Ikawa, Jean-Ju Chung

## Abstract

In mammalian sperm cells, regulation of spatiotemporal Ca^2+^ signaling relies on the quadrilinear Ca^2+^ signaling nanodomains in the flagellar membrane. The sperm-specific, multi-subunit CatSper Ca^2+^ channel, which is crucial for sperm hyperactivated motility and male fertility, organizes the nanodomains. Here, we report CatSperτ, the *C2cd6-*encoded membrane-associating C2 domain protein, can independently migrate to the flagella and serve as a major targeting component of the CatSper channel complex. CatSperτ loss-of-function in mice demonstrates that it is essential for sperm hyperactivated motility and male fertility. CatSperτ targets the CatSper channel into the quadrilinear nanodomains in the flagella of developing spermatids, whereas it is dispensable for functional channel assembly. CatSperτ interacts with ciliary trafficking machinery in a C2-dependent manner. These findings provide insights into the CatSper channel trafficking to the Ca^2+^ signaling nanodomains and the shared molecular mechanisms of ciliary and flagellar membrane targeting.

**Highlights:**

- CatSperτ encoded by *C2cd6* is a C2 membrane-associating domain containing protein
- CatSperτ loss-of-function impairs sperm hyperactivation and male fertility
- CatSperτ adopts ciliary trafficking machineries for flagellar targeting via C2 domain
- CatSperτ targets the CatSper channel into nanodomains of developing sperm flagella

## Introduction

Compartmentalization of the plasma membrane into distinct domains is an important mechanism to control signaling. Cilia and flagella are microtubule-based projections generated from basal bodies and ensheathed by the specialized membrane of distinctive lipid compositions (Garcia III et al., 2018; Walters et al., 2020). A conserved intraflagellar transporter (IFT) system assembles the axoneme - the core structure common to both cilia and flagella - whereas ciliary and flagellar membranes contain concentrated ion channels and membrane receptors for signal transduction (Pablo et al., 2017; Trötschel et al., 2020; Wang et al., 2021). Over the past decades, our increased knowledge of ciliary targeting of membrane proteins has established that vesicular transportation and IFTs enable the cargoes to pass transition zone and membrane diffusion barrier (Nachury and Mick, 2019; Nachury et al., 2010). Cilia function as sensory antenna are crucial for diverse biological processes including phototransduction, olfaction, and Hedgehog signaling.

As specialized cilia, flagella of mammalian sperm have one particular task. The flagellum provides propulsion for the sperm cells to reach and fertilize the eggs in the female reproductive tract. For this arduous journey, flagellar membrane proteins also play essential roles in signal transduction to regulate sperm motility, highlighting their importance in male fertility (Wang et al., 2021). However, how flagellar membrane assembly is coordinated with other early spermiogenesis process remains largely unknown. Just like cilium biogenesis, IFT systems are predicted to deliver flagellar membrane proteins. Yet, the absence of IFT components compromises axonemal growth and results in very short or no flagella (Avidor-Reiss and Leroux, 2015), making it difficult to directly test this idea. It remains unclear whether flagellar membrane targeting uses the same ciliary membrane targeting machinery or distinct motors, and how it is achieved.

In mammalian sperm cells, spatiotemporal regulation of Ca^2+^ signaling relies on the quadrilinear organization of the Ca^2+^ signaling nanodomains in the flagellar membrane (Chung et al., 2014; Hwang et al., 2019), where the sperm-specific, multi-subunit CatSper Ca^2+^ channel complexes assemble into zigzag rows in each quadrant (Zhao et al., 2021). Previous studies have found that only spermatozoa with four intact CatSper nanodomains can develop hyperactivated motility, undergo acrosome reaction, and successfully arrive at the fertilizing site (Chung et al., 2017; Chung et al., 2014; Ded et al., 2020). Thus, flagellar targeting of the CatSper channel and its localization into the quadrilinear nanodomains is crucial for sperm motility and male fertility. However, its complexity in composition (Hwang et al., 2019; Lin et al., 2021; Zhao et al., 2021), the inseparable loss-of-function phenotypes of each transmembrane (TM) subunit in mice (Chung et al., 2011; Hwang et al., 2019; Qi et al., 2007), and the lack of heterologous systems for functional reconstitution have limited our understanding of the mechanism of the CatSper assembly and delivery to flagella.

Here, we report CatSperτ (tau, *<u>CatSper T</u>argeting subunit*) containing C2 membrane-associating domain encoded by *C2CD6* is critical for CatSper flagellar targeting and trafficking into the quadrilinear nanodomains. CatSperτ loss-of-function studies in mice reveal that CatSperτ targets the preassembled CatSper complexes to elongating flagella, where CatSperτ links the channel-carrying vesicles and motor proteins. C2 domain truncation of CatSperτ is sufficient to compromise the spatiotemporal trafficking of the CatSper channel into the nanodomains in developing spermatids and during sperm maturation. We demonstrate that mutant sperm still form the CatSper channel, albeit significantly decreased, and conduct calcium, but they fail to hyperactivate, rendering the male infertile. These findings provide new insight into the molecular mechanisms of flagellar targeting of membrane proteins in general and organizing the CatSper Ca^2+^ signaling nanodomains in mammalian spermatozoa.

## Results

### CatSperτ is a New CatSper Component with the Membrane-Associating C2 Domain

The unique quadrilinear CatSper nanodomains within flagellar membrane suggest that the CatSper channel complex might require a membrane-targeting molecule specialized for this task. C2CD6 (previously known as ALS2CR11) was identified as one of the proteins significantly reduced in *CatSper1*-null spermatozoa that lack the entire CatSper channel complex (Hwang et al., 2019; Zhao et al., 2021). The testis-specific *C2CD6* gene encodes two isoforms: both long (>200 kDa) and short (∼70 kDa) forms contain the conserved membrane-associating C2 domain in both mouse and humans (Figures 1A and 1B). Thus, we hypothesize that C2CD6 is a new CatSper component that mediates CatSper trafficking to the flagellar membrane and/or spatial partitioning to the nanodomains.

**Figure 1.**
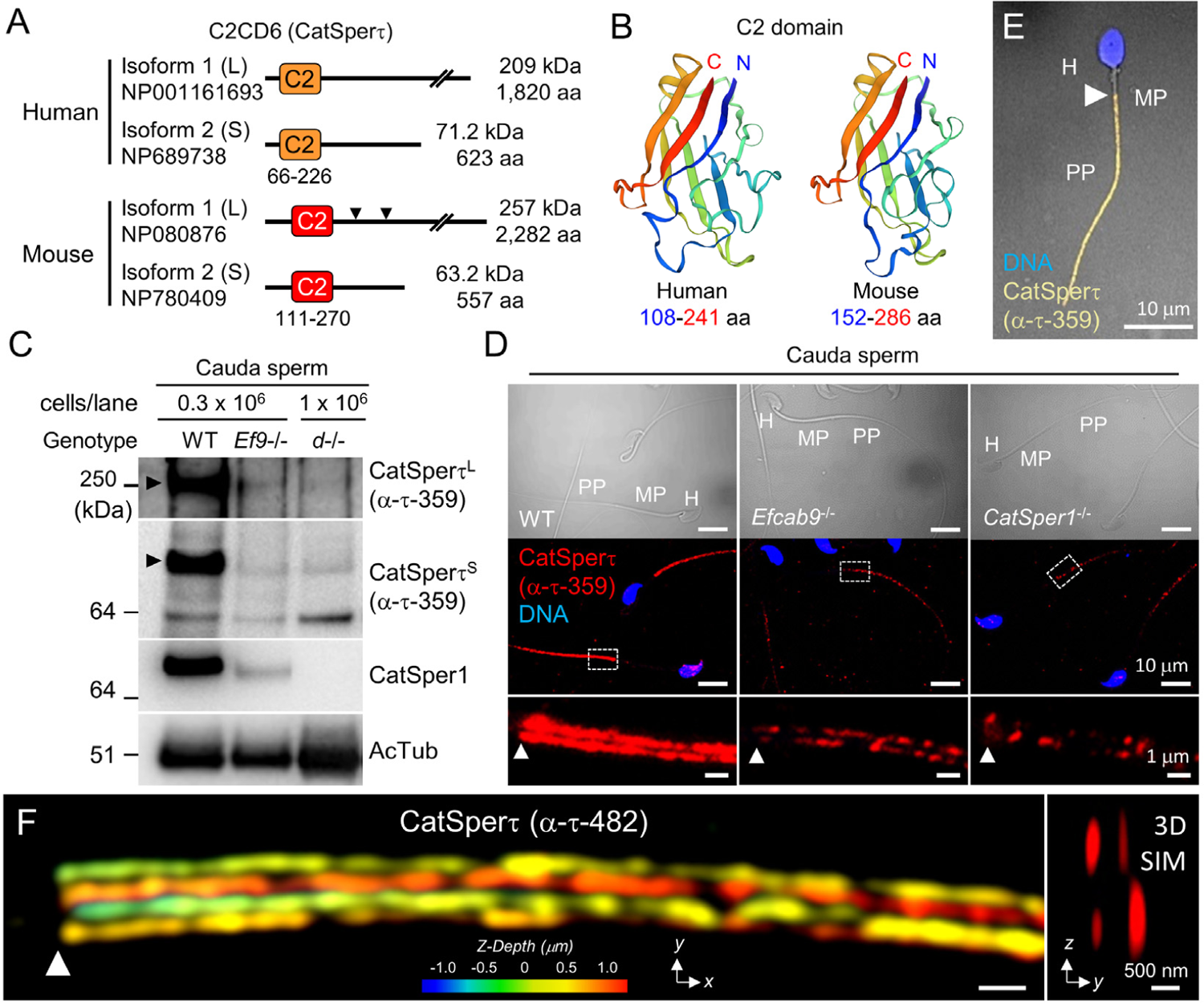
CatSpert is a new CatSper-associated C2 domain containing protein encoded by *C2cd6*. (A) Diagrams of human and mouse C2CD6 (CatSperτ) isoforms. C2 domain in human CatSperτ (amino acid (aa) position 66 – 226) and a counterpart region in mouse CatSperτ (aa position 111 – 270) are represented in orange and red boxes, respectively. Arrowheads indicate epitope regions of two CatSperτ antibodies (α-τ-359 and α-τ-482) used in this study. (B) Predicted structures of human (*left*) and mouse (*right*) CatSperτ C2 domains. N- and C-termini are colored in blue and red, respectively. (C-D) Association of CatSperτ with the CatSper channel in mouse sperm from cauda epididymis. (C) Immunoblotting of CatSperτ in *Efcab9*-null (*Ef9-/-*) and *CatSperd*-null (*d-/-*) sperm. Arrowheads indicate long (CatSperτ^L^) and short (CatSperτ^S^) isoforms. Acetylated tubulin (AcTub) is probed as a loading control. (D) Confocal images of immunostained CatSperτ in WT (*left*), *Efcab9*-null (*middle*), and *CatSper1*-null (*right*) epididymal sperm. DIC Images (*top*) and the corresponding fluorescence (*middle*) are shown. Each inset area in the fluorescence images is magnified (*bottom*). (E) A confocal image of immunostained CatSperτ in human sperm from ejaculate. Shown is a merged view of fluorescence and corresponding DIC images. Hoechst was used for DNA counterstaining. H, head; MP, mid-piece; PP, principal piece (D and E). (F) Quadrilinear arrangement of CatSperτ in WT mouse cauda sperm visualized by 3D structural illumination microscope (SIM) imaging. *z*-axis information is color-coded in *x*-*y* projection (*left*) and *y*-*z* projection (cross-section) is shown (*right*). Arrowheads indicate annulus, the junction between the midpiece and the principal piece (D-F). See also Figure S1.

We first examined whether C2CD6 expression is CatSper-dependent (Figures 1C-1F). Immunoblot analysis of sperm from a mouse cauda epididymis showed that protein levels of both long and short forms of C2CD6 are severely reduced in *CatSper1*- and *CatSperd*-null sperm that lack the entire CatSper channel, compared with those in WT sperm (Figures 1C and 1D, *right*). C2CD6 is localized in the principal piece in both mouse and human sperm (Figures 1D and 1E). 3D structured illumination microscopy (SIM) further revealed a quadrilateral arrangement of C2CD6 (Figure 1F), a hallmark of the CatSper Ca^2+^ signaling domain (Chung et al., 2017). In addition, the decreased C2CD6 proteins distribute discontinuously (Figures 1C and 1D, *middle*) in *Efcab9*-null sperm that contain overall only ∼ 20% of the protein levels of CatSper subunits and exhibit a fragmented pattern of the linear CatSper nanodomains (Hwang et al., 2019). All these results suggest that C2CD6 is associated with the CatSper channel complex in cauda sperm. C2CD6 is a newly identified *bona fide* CatSper component – we herein name it CatSperτ. Intriguingly, in *CatSper1*- or *CatSperd*-null sperm, CatSperτ is not completely absent, but still detected despite having even lower the protein levels compared to those of *Efcab9*-null sperm (Figures 1C and 1D). This minor yet obvious presence of CatSperτ in the absence of the channel complex distinguishes it from all the other previously reported CatSper subunits.

### CatSperτ Loss-of-Function Causes Male Infertility with Defective Sperm Hyperactivation

This expression pattern of CatSperτ further supports our hypothesis that CatSperτ regulates flagellar localization of the CatSper channel. To test this idea, we generated *CatSpert* mouse models by CRISPR/Cas9 genome editing (Figures 2 and S1). By injecting two pairs of guide RNAs - one targeting the first and 13^th^ exons and the other for the first and second exons - we obtained a knockout line that deletes 43.2 kb encoding almost the entire genomic region of CatSperτ^S^ and two mutant *CatSpert* lines that delete 134 bp (*134del*) or 128 bp (*128del*) of the protein coding region (Figures S1B-S1E). The mutant alleles were predicted to express mRNAs that encode frame-shifted proteins with early termination (Figure S1F). Our initial characterization found that homozygous *CatSpert-128del* and *CatSpert-134del* mice showed the identical phenotypes. *CatSpert-134del* line was used as *CatSpert*-mutant mice throughout this study unless indicated. We examined the protein levels of CatSperτ in mutant and knockout models in testes from WT, homozygous *CatSpert*-mutant (*CatSpert*^Δ/Δ^) and knockout (*CatSpert*^-/-^) males (Figures 2A and S1G). CatSperτ, mainly enriched in the microsome fraction, is absent from *CatSpert*^-/-^ testes (Figure S1G) whereas a few proteins with different molecular weights are distinguished from *CatSpert*^Δ/Δ^ testes by CatSperτ antibodies (Figures 2A and S1G). To characterize the proteins expressed specifically in *CatSpert*^Δ/Δ^ testes, we examined *CatSpert* mRNA levels in *CatSpert*^Δ/Δ^ testes (Figures S1H-S1L). In *CatSpert*^Δ/Δ^ testis, ∼40% of the *CatSpert* transcript is expressed compared with that of WT testes (Figure S1I). RT-PCR and Sanger sequencing demonstrated that *CatSpert*^Δ/Δ^ testes express truncated *CatSpert* mRNAs with 134 bp deletion or 186 bp deletion lacking exon 3 additionally (Figures S1J and S1K). These mRNAs are expected to generate mutant CatSperτ proteins with partial deletions in C2 domain at its N-terminus (Figure S1L). However, the mutant CatSperτ proteins were not detected sperm from *CatSpert*^Δ/Δ^ (i.e., *CatSpert*^Δ/Δ^ sperm; Figures 2B, 2C, and S2A, S2B), demonstrating that the C2-domain truncated CatSperτ is expressed in testes but fails to traffic to the *CatSpert*^Δ/Δ^ sperm flagella.

**Figure 2.**
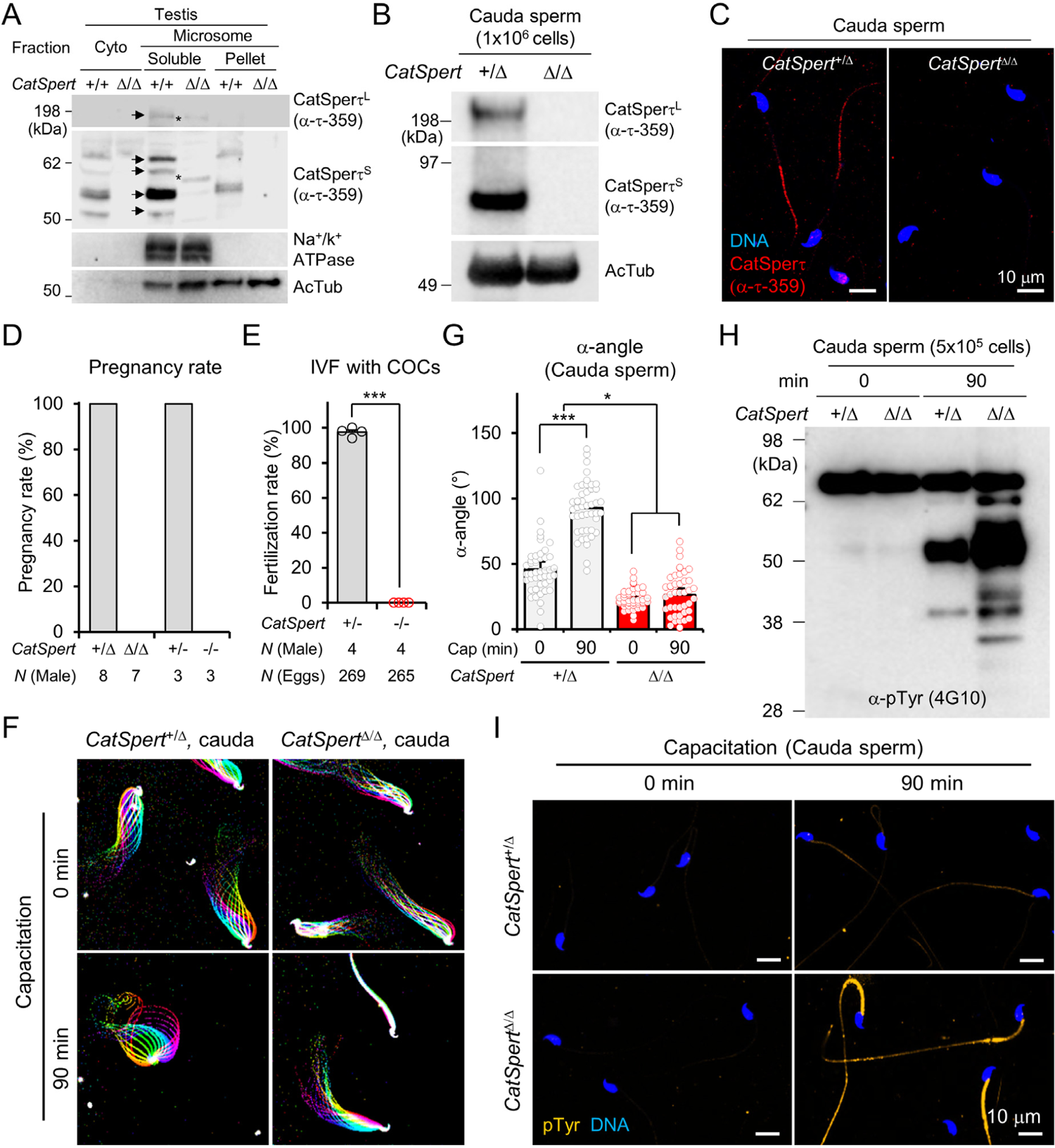
Genetic disruption of *CatSpert* impairs male fertility and sperm hyperactivation. (A-C) Validation of *CatSpert-*mutant mice generated in this study. CatSperτ protein expression was examined from the testis (A) and the cauda sperm (B and C) of homozygous *CatSpert-*mutant males by immunoblotting (A and B) and immunostaining (C). Arrows and asterisks indicate normal and mutant CatSperτ, respectively. Na^+^/K^+^ ATPase and acetylated tubulin (AcTub) were probed as loading controls (A and B). WT testes (A) and sperm from heterozygous *CatSpert*-mutant males (+/Δ) were used for positive control (B and C). (D) Percent pregnancy rates of fertile females mated with *CatSpert*-mutant (+/Δ and Δ/Δ) and knockout (+/- and -/-) males. Pregnancy rates of females mated with *CatSpert*-*134del* (+/Δ, N=5; Δ/Δ, N=4) and *128del* (+/Δ, N=3; Δ/Δ, N=3) are combined. (E) *In vitro* fertilization rates of *CatSpert*^+/-^ (N=4, 97.5 ± 1.3%) and *CatSpert^-/-^* (N=4, 0.0 ± 0.0%) sperm with cumulus oocyte complexes (COCs). Circles indicate *in vitro* fertilization rates of individual males. \*\*\**p*<0.001. (F) Flagella waveforms of *CatSpert*^+/Δ^ (*left*) and *CatSpert*^Δ/Δ^ (*right*) sperm from cauda epididymis. Tail movements of the sperm tethered to imaging chamber were recorded at 200 fps speed before (0 min, *top*) and after (90 min, *bottom*) inducing capacitation *in vitro*. Overlays of flagellar waveforms for two beat cycles are color-coded in time. (G) Maximum angles of the primary curvature in the midpiece (α-angle) of cauda sperm. Heterozygous (+/Δ, gray bars) and homozygous (Δ/Δ, red bars) *CatSpert*-mutant sperm before (0min; +/Δ, 45.76 ± 2.96°; Δ/Δ, 23.28 ± 1.16°) and after (90 min; +/Δ, 92.71 ± 3.02°; Δ/Δ, 26.13 ± 2.51°) inducing capacitation (Cap). Tail parallel to head is set to 0°. Circles indicate the measured α-angle of the individual sperm cells (N=45) from 3 males in each group. \**p*<0.05 and \*\*\**p*<0.001. (H-I) Excessive development of protein tyrosine phosphorylation (pTyr) in *CatSpert*^Δ/Δ^ cauda sperm during capacitation. (H) Immunoblotting of protein tyrosine phosphorylation in *CatSpert*-mutant sperm cells before and after capacitation (0 and 90 min, respectively). (I) Confocal images of pTyr-immunostained *CatSpert-*mutant sperm cells. Shown are images of *CatSpert*^+/Δ^ (*top*) and *CatSpert*^Δ/Δ^ (*bottom*) sperm before (*left*, 0 min) and after (*right*, 90 min) inducing capacitation. Data represented as mean ± SEM (E and G). Hoechst to counterstain DNA (C and I). See also Figures S1 and S2 and Video S1.

Both homozygous *CatSpert-*mutant and knockout mice have no gross abnormality in survival, appearance, or behavior. *CatSpert*^Δ/Δ^ and *Catspert*^-/-^ females have normal reproductive ability and give birth with no difference in litter size. However, both *CatSpert*^Δ/Δ^ and *CatSpert*^-/-^ males are infertile (Figures 2D, 2E, and S2C), despite normal testis histology, sperm morphology, and sperm counts in cauda epididymis (Figures 2C and S2A, S2D, and S2E). Although the *CatSpert*^Δ/Δ^ males express mutant CatSperτ containing truncated C2 domain in testis, they show identical physiological characteristics to those of *CatSpert*^-/-^ males. The shared phenotypes between *CatSpert*^Δ/Δ^ and *CatSpert*^-/-^ mice highlight that the C2 domain is essential for CatSperτ function. We noted that *CatSpert*^Δ/Δ^ mice are not only suitable for CatSperτ loss-of-function in mature cauda sperm but also ideal to investigate the critical N-terminal subdomain of C2 in the native context during male germ cell development. Therefore, we mainly used the *CatSpert*-mutant mice in this study.

To understand how CatSperτ loss-of-function causes male infertility, we analyzed sperm motility and their flagellar movement in *CatSpert*^Δ/Δ^ males (Figures 2F, 2G, and S2F-S2H). Computer-assisted semen analysis (CASA) revealed that swimming velocities (i.e., VCL and VAP) and lateral head displacement were significantly lower in *CatSpert*^Δ/Δ^ sperm than those in sperm from *CatSpert^+^*^/Δ^ males (i.e., *CatSpert^+^*^/Δ^ sperm) after incubating them under capacitation condition (Figure S2F). *CatSpert*^Δ/Δ^ sperm also failed to increase the maximum angle of midpiece curvature (α-angle) and to reduce tail beating speed (Figures 2F, 2G, and S2G, S2H; Video 1), other known features of hyperactivation (Qi et al., 2007). All the results demonstrate that CatSperτ loss-of-function impairs hyperactivated motility and causes male infertility.

### CatSperτ-Deficiency Reduces the CatSper Protein Levels and Disorganizes the Nanodomains

Under capacitating conditions, a complete lack of or insufficient Ca^2+^ entry prevents sperm from developing hyperactivated motility, controlling PKA activity, and developing the subsequent protein tyrosine phosphorylation (pTyr) (Chung et al., 2017; Chung et al., 2014; Navarrete et al., 2015). *CatSpert*^Δ/Δ^ sperm prematurely potentiate capacitation-associated PKA activity (Figure S2I) and pTyr (Figures 2H and 2I) compared to *CatSpert*^+/Δ^ sperm, suggesting a compromised Ca^2+^ influx. Thus, we hypothesized that CatSperτ deficiency dysregulates the level of CatSper channel and/or its function in sperm. We examined CatSper TM (CatSper1-4, β and δ) and non-TM (CatSperζ and EFCAB9) subunit levels and found that they are only ∼10% of mature *CatSpert*^Δ/Δ^ sperm from cauda epididymis compared to those of WT sperm (Figures 3A and 3B). Interestingly, we found that the CatSper channel, probed by α-CatSper1, is localized at the distal region of the principal piece in the *CatSpert*^Δ/Δ^ sperm (Figures 3C and S3A, S3B). 3D SIM imaging further suggested that the CatSper channel lacking CatSperτ does not exhibit the typical quadrilateral nor continuous linear distribution of the nanodomains (Figure 3D). Based on these results, we propose that CatSperτ might play a crucial role in linear and quadrilateral distribution of the channel complexes in the flagellar membraneduring tail formation and/or epididymal maturation.

**Figure 3.**
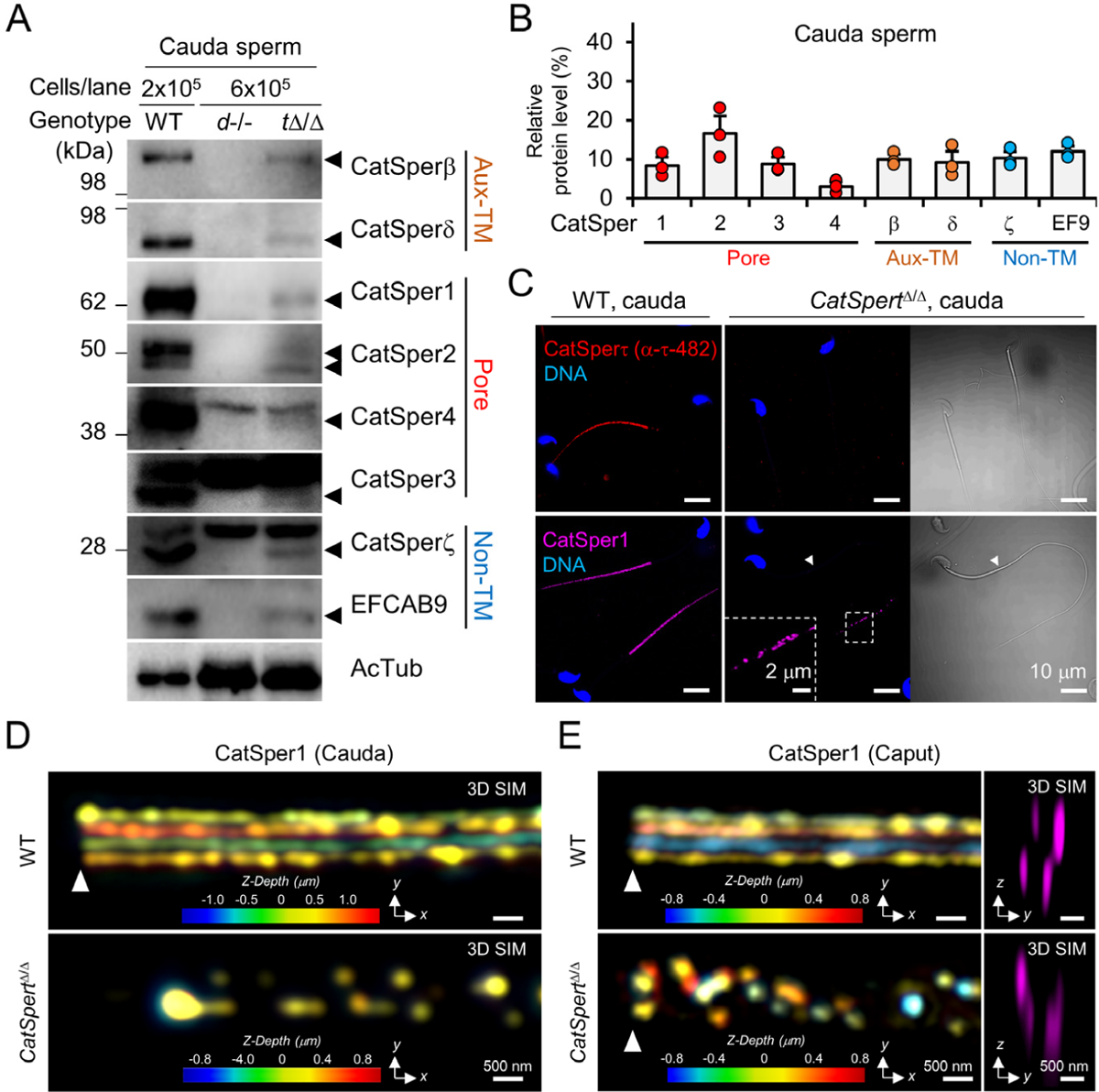
CatSpert loss-of-function diminishes CatSper subunits and disorganizes the CatSper nanodomain in epididymal sperm. (A-B) Reduced protein levels of CatSper subunits in *CatSpert*^Δ/Δ^ sperm from cauda epididymis. (A) Immunoblotting of CatSper subunits in *CatSpert*^Δ/Δ^ cauda sperm. Arrowheads indicate the bands of the target subunits (Aux-TM: auxiliary transmembrane, Pore: pore-forming, and Non-TM: non-transmembrane). *CatSperd*-null sperm (*d*-/-) is a negative control for the absence of the whole channel. Acetylated tubulin (AcTub) is a loading control. (B) Relative protein levels of the CatSper subunits in *CatSpert*^Δ/Δ^ cauda sperm. Around 10% of each subunit is detected in *CatSpert*^Δ/Δ^ sperm compared to WT sperm; CatSper1 (8.4 ± 2.1%), 2 (16.6 ± 4.5%), 3 (8.8 ± 1.7%), 4 (3.0 ± 1.2%), β (10.0 ± 1.2%), δ (9.3 ± 2.8%), ζ (10.3 ± 1.7%) and EFCAB9 (EF9, 12.0 ± 1.4%). Circles indicate relative levels of CatSper subunits in *CatSpert*^Δ/Δ^ sperm from individual males. Protein levels were quantified by measuring the band density from the independent western blots. Data represented as mean ± SEM. N=3. (C) Diminished CatSper signals detected at the distal flagella in *CatSpert*^Δ/Δ^ sperm. Shown are confocal images of immunostained CatSperτ (*top*) and CatSper1 (*bottom*) in WT (*left*) and *CatSpert*^Δ/Δ^ (*middle* and *right*) cauda sperm and the corresponding DIC images (*right*). Sperm heads were counterstained with Hoechst. A magnified inset is shown (*bottom middle*). (D-E) 3D SIM images of CatSper1 in cauda (D) and caput (E) epididymal sperm from WT (*top*) and *CatSpert*^Δ/Δ^ (*bottom*) males. Colors in *x*-*y* projections encode *z*-depth distance from the focal plan (D and E, *left*). *y*-*z* cross sections are shown at right (E). Arrowheads indicate annulus (C, D, and E). See also Figure S3.

To test this idea, we first examined the protein levels of CatSper subunits and the nanodomain organization of WT and *CatSpert*^Δ/Δ^ sperm from the caput and corpus epididymis (Figures 3E and S3C-S3G). Corpus epididymal sperm from *CatSpert*^Δ/Δ^ males express around 10 - 25% of CatSper subunits compared to those in WT (Figures S3C and S3D). WT sperm from the caput epididymis displayed four CatSper nanodomains (Figure 3E, *top*), indicating the linear Ca^2+^ signaling domains are organized during spermiogenesis in testis. In *CatSpert*^Δ/Δ^ caput sperm, however, we noted a few striking differences. First, the CatSper channel presented only 2 or 3 discontinuous linear nanodomains (Figure 3E, *bottom*), suggesting its role in organizing and maintaining the nanodomains. Second, regardless of the low protein levels, the mutant CatSper channel is still detected in the proximal principal piece, as in the WT channel of cauda sperm, suggesting changes in CatSper channel distribution in *CatSpert*^Δ/Δ^ sperm during epididymal maturation. Indeed, the CatSper channels showed transitions in their localization from the proximal to distal region of the flagella in *CatSpert*^Δ/Δ^ sperm and diminution of the protein levels (Figures S3D, S3F and S3G).

### CatSperτ Is Dispensable for Functional CatSper Channel Assembly

Delineation of the CatSperτ function in flagellar trafficking is complicated by the significantly low protein levels of the CatSper channel in *CatSpert*^Δ/Δ^ sperm flagella (Figures 3 and S3). To better understand why the protein levels are reduced, we first tested whether the functional CatSper channel is assembled in epididymal *CatSpert*^Δ/Δ^ sperm.

Testis co-immunoprecipitation (coIP) showed that CatSperτ is in complex with CatSper1 and CatSperδ in WT testis (Figure 4A). However, we found that all examined CatSper subunits could form immunocomplexes with CatSper1 and CatSperδ in both *CatSpert*^Δ/Δ^ (Figures 4B and S4A) and *CatSpert*^-/-^ (Figures 4C and S4B) testes. These results suggest CatSperτ is dispensable in forming the CatSper channel complex in testes (Figures 4B, 4C, and S4A, S4B).

**Figure 4.**
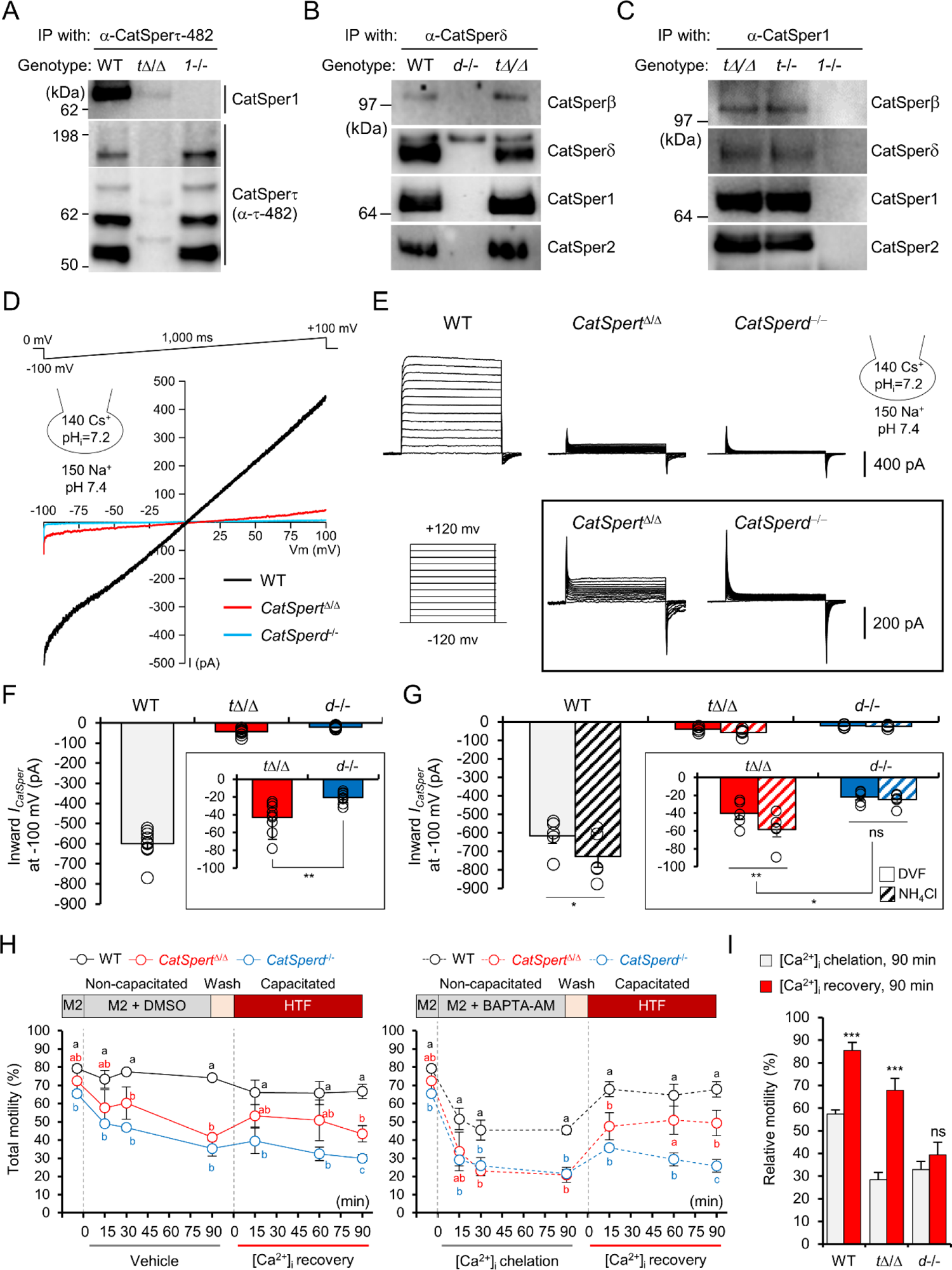
Functional CatSper channel is assembled in the absence of CatSpert. (A) CatSperτ in complex with CatSper channel in testis. (B-C) CatSper subunits in complex without CatSperτ in testes. Pore (Catsper1 and 2) and auxiliary transmembrane subunits (CatSperβ and δ) were detected from CatSperδ-(B) and CatSper1-(C) immunocomplexes from solubilized testis microsome. (D-E) Representative traces of CatSper current (*I_CatSper_*) from WT, *CatSpert*^Δ/Δ^, and *CatSperd*^-/-^ corpus sperm. *I_CatSper_* was elicited by voltage ramp (−100 to + 100 mV) from 0 mV holding potential (D) or by step protocol (−120 to +120 mV) in 20 mV increments (E). The cartoons represent pH and monovalent ion composition in pipet and bath solutions (D and E). (F) Inward *I_CatSper_* measured from WT (gray, N=9), *CatSpert*^Δ/Δ^ (red, N=9), and *CatSperd*^-/-^ (blue, N=8) corpus sperm at −100 mV. (G) Increase of inward *I_CatSper_* by intracellular alkalization. The currents before (solid) and after (hatched) adding 10 mM NH_4_Cl to bath solution were recorded at −100 mV from WT (gray), *CatSpert*^Δ/Δ^ (red), and *CatSperd*^-/-^ (blue) corpus sperm (N=5, each). Bars and circles indicate the average *I_CatSper_* and the currents from individual sperm, respectively (F and G). Insets show magnified *I_CatSper_* traces in *CatSpert*^Δ/Δ^ and *CatSperd*^-/-^ sperm (F and G). (H-I) Effect of intracellular Ca^2+^ (Ca^2+^_i_) changes on motility of WT, *CatSpert*^Δ/Δ^, and *CatSperd*^-/-^ cauda sperm. (H) Time-course changes on total sperm motility. Motility was compared from vehicle (*left*) or BAPTA-AM (*right*) treated WT (black), *CatSpert*^Δ/Δ^ (red), and *CatSperd*^-/-^ (blue) sperm. Different letters indicate significant differences between the genotypes at each time point. (I) Relative motility changes after Ca^2+^_i_ chelation and recovery. Motility at 90 min time points from Ca^2+^_i_ chelation (grey) and recovery (red) were normalized with the motility before loading BAPTA-AM (M2 in H) in each genotype. Relative motility after Ca^2+^_i_ chelation and recovery within each genotype was compared. N=3. ns, non-significant, *\*p*<0.05, *\*\*p*<0.01, and *\*\*\*p*<0.001 (F, G, and I). Data represented as mean ± SEM (F, G, H, and I). See also Figure S4.

To directly test whether the functional CatSper channel is assembled without CatSperτ, we measured CatSper current (*I_CatSper_*) by electrophysiological recording (Figures 4D-4G). The inward current from *CatSpert*^Δ/Δ^ sperm is only ∼6% of the current from WT sperm but still significantly larger than that of *CatSperd*-null sperm (Figures 4D-4F), which lack the entire CatSper channel (Chung et al., 2011). Furthermore, adding 10 mM NH_4_Cl increased *I_CatSper_* in *CatSpert*^Δ/Δ^ sperm (Figure 4G), consistent with alkalinization activation of CatSper (Kirichok et al., 2006). Thus, *CatSpert*^Δ/Δ^ sperm assembled the functional CatSper channel with the previously known key characteristics despite the ∼10% protein levels of CatSper subunits (Figures 3 and S3).

Intriguingly, Ca^2+^ conductance in *CatSpert*^Δ/Δ^ sperm was not enough to develop hyperactivated motility in *CatSpert*^Δ/Δ^ sperm (Figures 2 and S2). In agreement, we found that the reduced motility of *CatSpert*^Δ/Δ^ sperm either by extended incubation or intracellular Ca^2+^ chelation can be only partially rescued (Figures 4H and 4I), suggesting limited CatSper-mediated Ca^2+^ entry during recovery or capacitation. In particular, *CatSpert*^Δ/Δ^ sperm fail to fully recover their curvilinear velocity (VCL) compared to the initial velocity after inducing capacitation (Figures S4C and S4D). This is in stark contrast to *Efcab9^-/-^* sperm, which present ∼30% proteins levels of CatSper subunits and ∼60% Ca^2+^ conductance, were able to recover nearly the full extent of VCL and ALH (Chung et al., 2017; Hwang et al., 2019).

### CatSperτ Is Essential for CatSper Channel Targeting to Flagella in Developing Spermatids

Given that the CatSper channel lacking CatSperτ is functional (Figures 4 and S4), we next tested whether the CatSperτ loss-of-function compromises targeting of the assembled CatSper channel complex to the flagella in developing germ cells. Flagellated mouse spermatids undergo dramatic morphological changes during development (Clermont et al., 1993). Notably, the C2-truncated mutant CatSperτ did not affect the CatSper subunits to form complexes as early as in round spermatids (Figures 5A and 5B) isolated by STA-PUT (Miller and Phillips, 1969). As thin flagellum - without the outer dense fiber and the fibrous sheath - begins to protrude at this stage (Figure 5C), we hypothesized that the CatSperτ regulates the transport of the CatSper channel from the cell body to the developing tail.

**Figure 5.**
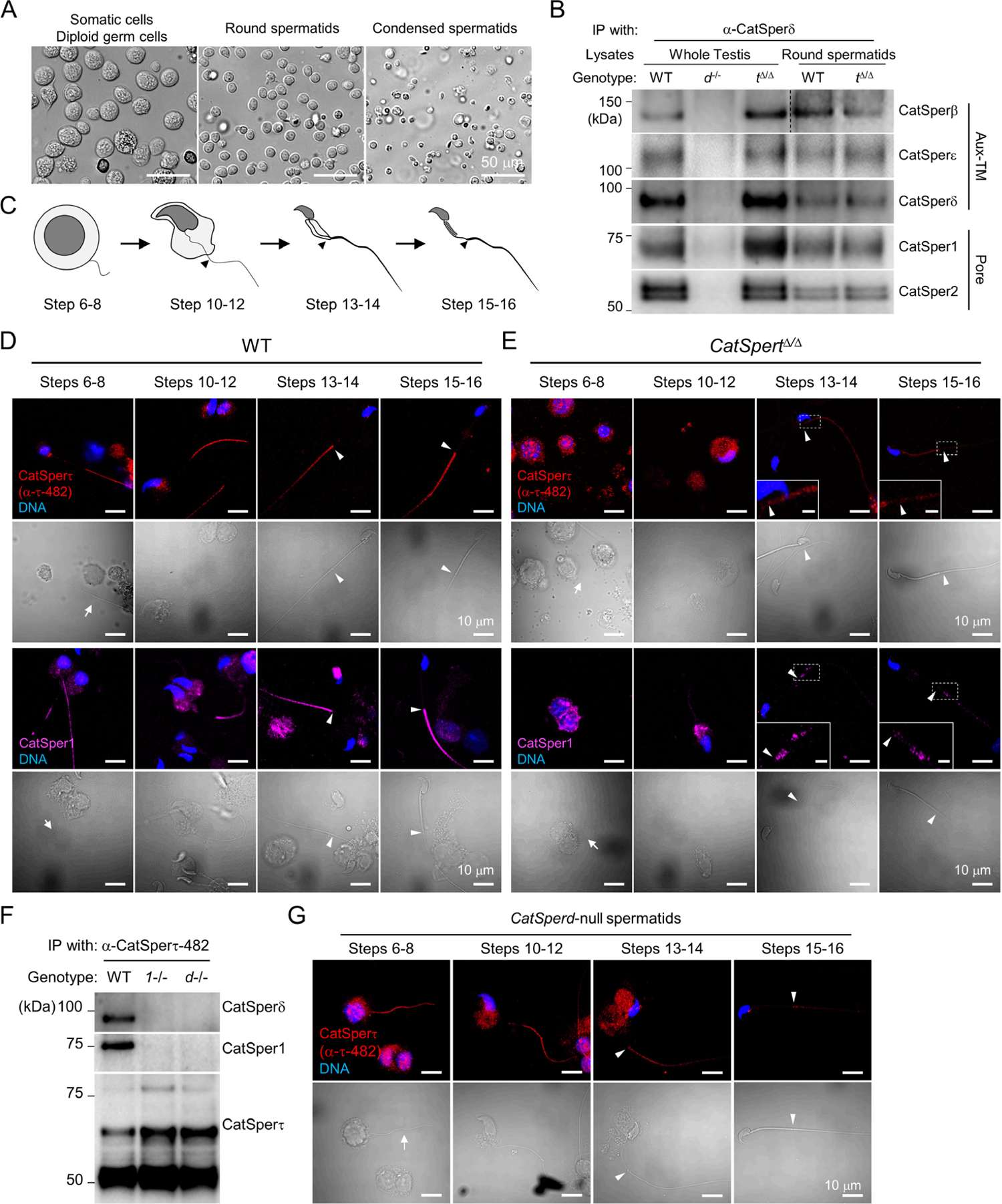
Loss of CatSpert function delays CatSper channel targeting to the flagella during sperm development. (A-B) Formation of the CatSper channel complex in WT and *CatSpert*^Δ/Δ^ round spermatids. (A) Separation of germ cell populations by STA-PUT. Fractions enriched in somatic and diploid germ cells (*left*), round spermatids (*middle*), and condensed spermatids (*right*) are shown. (B) Detection of CatSper pore (CatSper1 and 2) and auxiliary transmembrane subunits (Aux-TM; CatSperβ, ε, and δ) from the CatSper-immunocomplexes in both WT and *CatSpert*^Δ/Δ^ round spermatids (RS). WT and *CatSperd*-null (*d*-/-) testes lysates were used for positive and negative control, respectively. (C) Depiction of flagella development in elongating spermatids. Mouse spermatids from developmental steps from 6 to 16 were classified into four groups by morphological characteristics: steps 6-8 (spherical cell body and protruding axoneme), steps 10-12 (hook-shaped sperm head and elongating axoneme), steps13-14 (condensed sperm head and flagella with developed principal piece), and steps 15-16 (compartmentalized mitochondrial sheath at midpiece). (D-E) Confocal images of immunostained CatSperτ (*top*) and CatSper1 (*bottom*) in WT (D) and *CatSpert*^Δ/Δ^ (E) spermatids. Magnified insets show the proximal principal piece of *CatSpert*^Δ/Δ^ spermatids (E, scale bar = 2 μm). (F-G) CatSperτ trafficking to the prospective principal piece of the flagella in elongating spermatids in the absence of the CatSper complex. (F) Interaction of CatSperτ with CatSper1 or δ in WT, but not in *CatSperd*-null *(d*-/-) or *CatSper1*-null (*1*-/-) testis. (G) Confocal images of CatSperτ in *CatSperd*-null spermatids. CatSperτ transiently localizes to the flagella of *CatSperd*-null spermatids. Arrows and arrowheads indicate flagella in step 6-8 spermatids and annulus, respectively (D, E, and G). Hoechst to counterstain DNA (D, E, and G). See also Figure S5.

To test this hypothesis, we compared the intracellular localization of the CatSperτ and the CatSper channel, probed by α-CatSper1 or δ, in developing WT and *CatSpert*^Δ/Δ^ spermatids according to the developmental steps, well classified by the morphological characteristics (Figures 5C-5E and S5). We observed that the CatSperτ as well as the CatSper channel complex are localized at the prospective principal piece of WT flagella as early as steps 6-8 spermatids (Figures 5D and S5A). In *Catspert*^Δ/Δ^ spermatids, however, confocal imaging detected that the C2-domain truncated mutant CatSperτ, as well as CatSper1 and δ, is mostly enriched in the cell body until step 12 (Figure 5E and S5B). At steps 13-14 spermatids, a weak mutant CatSperτ signal was detected but the positive mutant signal eventually disappeared in later steps of mutant spermatids (Figure 5E) and epididymal sperm (Figures 2C and 3C), suggesting removal of the mutant proteins. By contrast, weak and punctate CatSper1 or δ signals observed in *CatSpert*^Δ/Δ^ spermatids from the steps 13-14 remain in the spermatids until at later steps (Figures 5E and S5B) and in the epididymal sperm (Figures 2C and 3C). All these results illustrate that C2 domain truncation compromises the flagellar targeting of CatSperτ and assembled CatSper channel in developing spermatids.

Small amount of CatSperτ remains in the cauda epididymal *CatSper1*-null sperm that lack the entire CatSper channel complex (Figure 1), suggesting CatSperτ could be an upstream molecule among the CatSper components for the CatSper targeting in developing spermatids. Thus, we tested whether CatSperτ can traffic to the flagella without associating with the assembled CatSper channel. In *CatSperd*- or *CatSper1*-null testis, CatSperτ was not found in the immunocomplex of the CatSper pore (CatSper1) or auxiliary TM (CatSperδ) subunits, respectively (Figure 5F). This result indicates that CatSperτ normally associates with the fully assembled channel complex; the small amount of CatSperτ detected in *CatSper1*-null sperm (Figure 1D) is presumably targeted to the flagella without associating with the channel complex. To further clarify that CatSperτ traffics to sperm flagella in absence of the CatSper channel, we observed the intracellular localization of CatSperτ in developing *CatSper1*-null and *CatSperd*-null spermatids. Contrary to the first appearance of the CatSper channels in *CatSpert*^Δ/Δ^ spermatids at steps 13-14 (Figure 5E), CatSperτ was observed in the flagella of *CatSper1*-null and *CatSperd*-null spermatids as early as the step 6 (Figures 5G and S5D) just like in WT spermatids (Figure 5D). The CatSperτ signal, however, becomes gradually weaker in the spermatids lacking CatSper channel complex at later stages (steps 13-16). These results demonstrate that CatSperτ serves as an upstream CatSper component in the CatSper flagellar targeting, which requires association with the channel complex to remain in the flagella after trafficking.

### CatSperτ Targets the Channel Complex to the Quadrilinear Nanodomains in the Flagella

CatSperτ loss-of-function dysregulates the flagellar targeting of the CatSper channel in developing spermatids (Figures 5 and S5), which results in impaired quadrilinear distribution of the CatSper channels in epididymal sperm flagella (Figures 3 and S3). Thus, we hypothesized that CatSperτ promotes quadrilinear distribution of the assembled CatSper channel in developing spermatids. To test this idea, we examined the nanodomain structures in developing spermatids by using 3D SIM imaging. CatSperτ and the assembled CatSper channel complex are all arranged quadrilinearly along the principal piece of WT spermatids at steps 13-16 (Figures 6A and S6A). In *CatSpert*^Δ/Δ^ spermatids, however, the weak signals of the mutant CatSperτ and CatSper channel are resolved as discontinuous nanodomains (Figure 6B). Intriguingly, we were able to further resolve the CatSperτ distribution in *CatSper1*- and *CatSperd*-null spermatids at steps 13-14 (Figures 6C and S6B) when the positive CatSperτ signal becomes weaker (Figures 5G and S5D). Despite discontinuity in the signals, CatSperτ clearly showed the typical quadrilateral distribution along the principal piece in *CatSper1*-null and *CatSperd*-null spermatids, supporting the idea that CatSperτ presumably contributes to the four-nanodomain formation.

**Figure 6.**
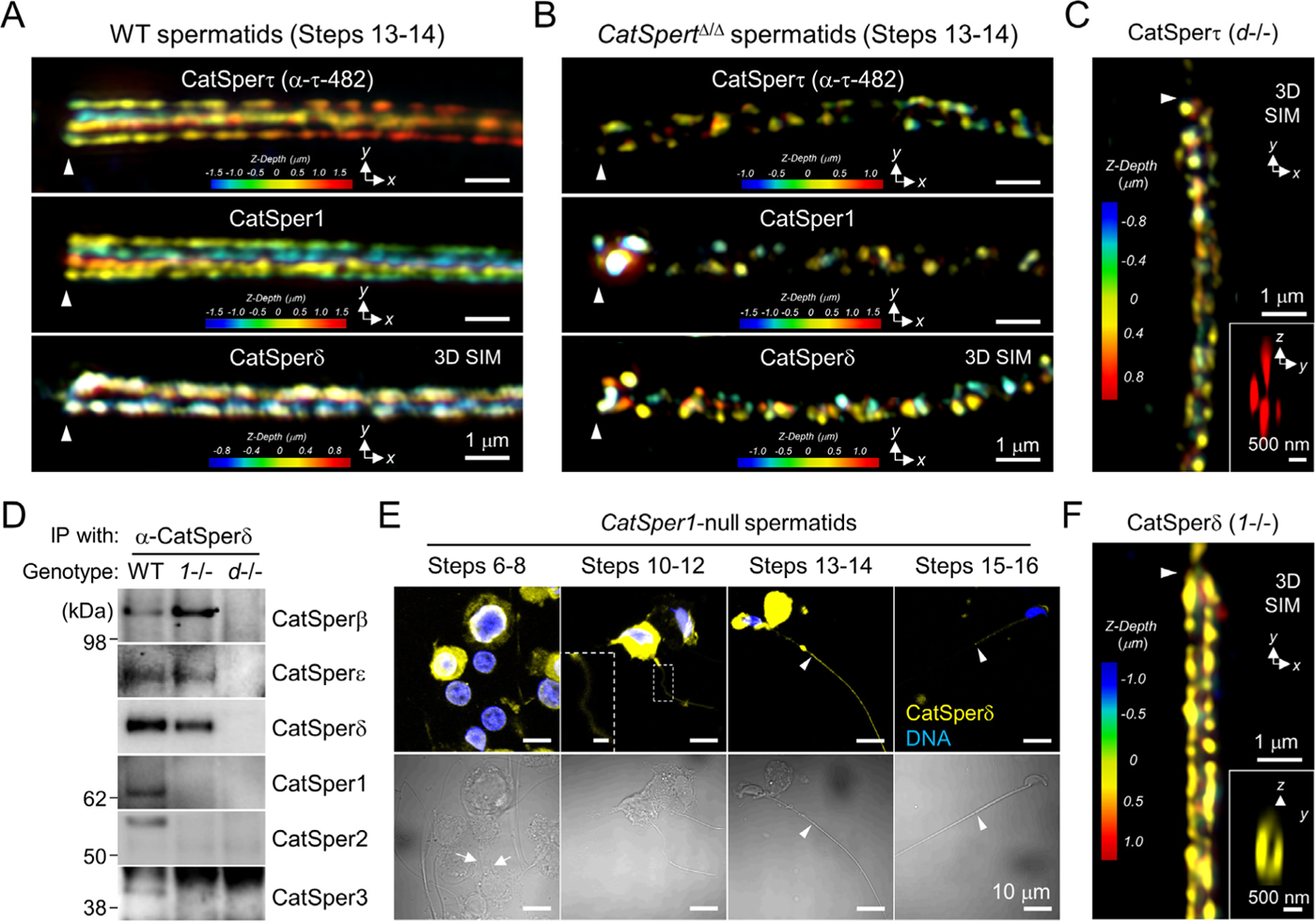
CatSpert is a major CatSper component to regulate the channel targeting to the nanodomains in elongating spermatids. (A-B) 3D SIM images of CatSperτ (*top*), CatSper1 (*middle*), and CatSperδ (*bottom*) in WT (A) and *CatSpert*^Δ/Δ^ (B) spermatids at steps13-14. (C) 3D SIM imaging of CatSperτ in *CatSperd*-null spermatid at step 13. An inset representing the *y*-*z* cross section image depicts quadrilateral arrangement. *z*-axis information was color-coded. α-CatSperτ-482 was used for A, B, and C. (D-F) Minimal contribution of the auxiliary TM subunits to flagellar trafficking in the absence of the tetrameric channel. (D) CatSper Aux-TM subunits (CatSperβ, δ, and ε) co-immunoprecipitated in *CatSper1*-null (*1*-/-) testis but not with CatSperτ. See also Figure 5F. WT and *CatSperd*-null testes were used for positive and negative control, respectively. (E) Confocal images of immunostained CatSperδ in *CatSper1*-null spermatids. Corresponding DIC images of immunostained spermatids are shown at bottom. A magnified inset (scale bar = 2 μm) shows faint CatSperδ signal in flagellum. Hoechst to counterstain DNA. (F) A 3D SIM image of CatSperδ in a step 13 *CatSper1*-null spermatid. *y*-*z* cross section shown at inset depicts bilateral localization. Information for *z*-axis was color-coded. Arrows and arrowheads indicate flagella (E) and annulus (A, B, C, E, and F), respectively. See also Figure S6.

It is of note that condensed *CatSpert*^Δ/Δ^ spermatids (Figures 5 and S5) and mature epididymal sperm (Figures 3 and S3) retain the small amount of the CatSper channel in the flagella, suggesting the possible involvement of another CatSper components in the CatSper channel targeting to sperm flagella, i.e. CatSperδ (Chung et al., 2011). Intriguingly, CatSperδ forms the complex that contains the auxiliary TM subunits alone but not the pore-forming subunits in the absence of CatSper1 (Chung et al., 2011; see also Figure 6D). Thus, we tested whether the auxiliary TM subunits are also involved in flagellar trafficking in developing spermatids. Confocal imaging revealed that the auxiliary TM subunits, probed by α-CatSperδ, show positive signal transiently at the prospective principal piece of *CatSper1*-null spermatids at steps 10-14 (Figure 6E). Given that CatSperδ did not immunocomplex with other pore subunits nor CatSperτ in *CatSper1*-null testis (Figures 5F and 6D), this result indicates that a complex comprised of the auxiliary TM subunits can traffic to flagella alone in developing spermatids just like CatSperτ. Closer observation by 3D SIM imaging, however, unraveled its bilateral distribution along the flagella of *CatSper1*-null spermatids at step 13 (Figure 6F). This result indicates that the CatSperδ-complex containing TM auxiliary subunits can traffic to the flagella but is not sufficient to distribute to the four linear nanodomains. By contrast, CatSper1 was not associated with other pore subunits in the absence of CatSperδ (Figure S6C) and fail to traffic to the flagella of developing spermatids from *CatSperd*-null males (Figure S6D). Thus, CatSper pore-forming subunits are presumably not able to traffic to flagella without other components. All these results from the developing spermatids and epididymal sperm from WT, *CatSper1*^-/-^, *CatSperd*^-/-^, and *CatSpert*^Δ/Δ^ males support that CatSperτ is a key component not only for flagellar targeting of the assembled CatSper channel but for their quadrilateral compartmentalization (Figure S6E).

### CatSperτ Mediates the CatSper Channel Localization via Cytoplasmic Vesicles and Motor Proteins

CatSperτ traffics to elongating flagella independent of CatSper channel complex in developing spermatids (Figures 6 and S6). To understand the molecular mechanisms of CatSperτ in flagellar targeting and to determine the effect of the C2 truncation in the subcellular localization, we expressed recombinant WT and C2-domain truncated mutant CatSperτ (Δ103-164; see also Figure S1) in the heterologous systems (Figure 7A). We found the lentiviral transduced WT CatSperτ was enriched as puncta at the ciliary base in ciliated hTERT-RPE1 (Figures 7B and S6F) and 293T (Figure S6G). Intriguingly, mutant CatSperτ was simply diffused throughout the cytoplasm and barely formed puncta. WT CatSperτ puncta were also observed from non-ciliated 293T (Figures S7A-C) and hTERT-RPE1 (Figure 7C) cells near the centrosome. This specific localization pattern of WT CatSperτ in the heterologous systems indicates CatSperτ can be recruited close to the centrosome originated basal body prior to the flagellar targeting. Furthermore, transiently expressed recombinant CatSper1 and CatSperζ were observed near the WT CatSperτ puncta (Figures 7D and S7D), suggesting that CatSperτ brought those CatSper subunits to the basal body. These results indicate that CatSperτ, when associated with the assembled CatSper channel complex, can modulate transportation of the channel to the basal body.

**Figure 7.**
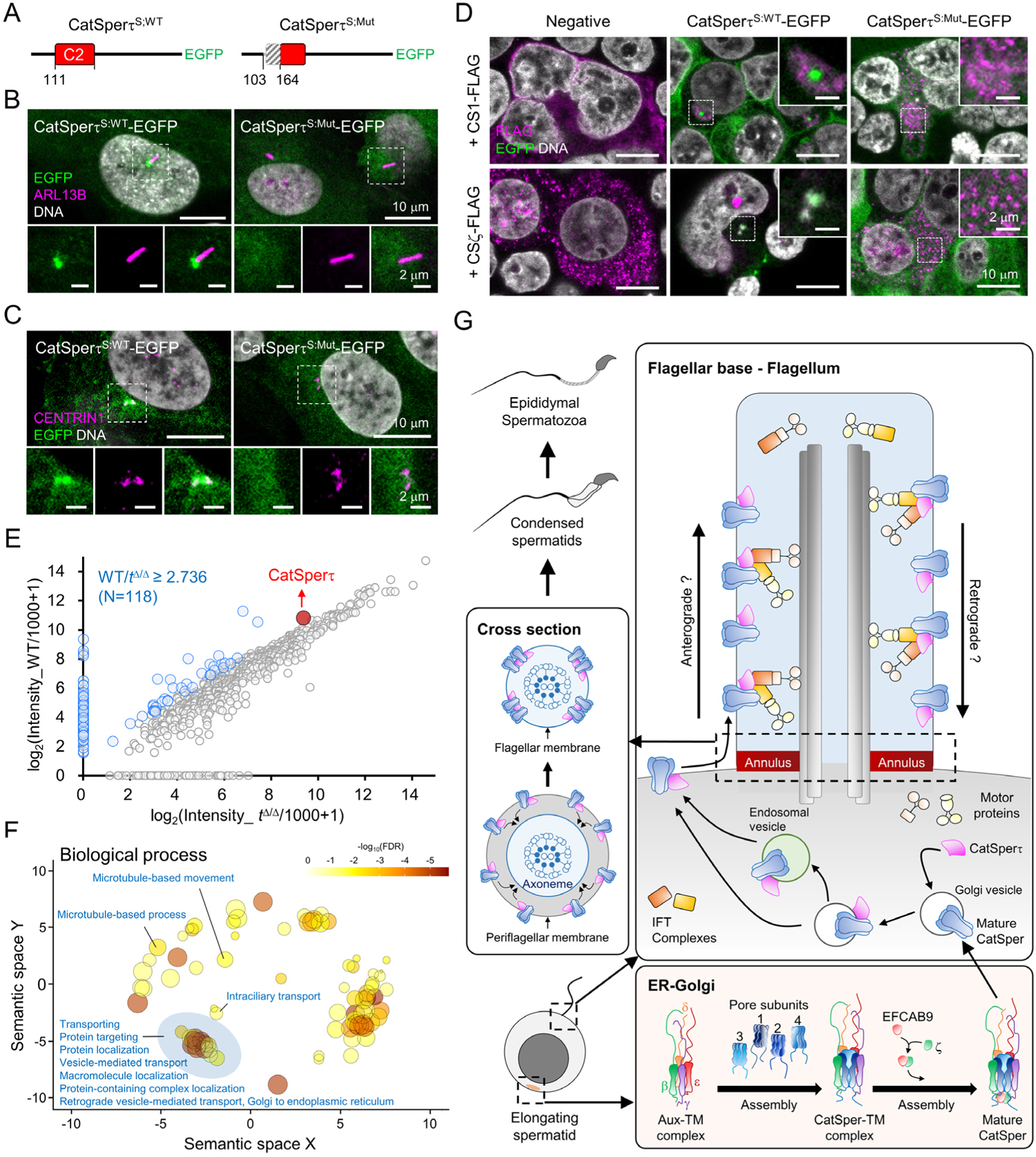
CatSperτ modulates the intracellular localization and flagellar targeting of the CatSper channel by interacting with cytoplasmic vesicles via C2 domain. (A) Recombinant short forms of WT (CatSperτ^S:WT^-EGFP) and C2-truncated mutant CatSperτ (CatSperτ^S:Mut^-EGFP) proteins. EGFP-tagged proteins were expressed by lentiviral transduction in heterologous system. (B-C) Confocal images of ciliated (B) and non-ciliated (C) hTERT-RPE1 cells stably expressing EGFP-tagged WT (*left*) and mutant (*right*) CatSperτ. ARL13B (B) and CENTRIN1 (C) were immunostained to label cilia and centrosome, respectively. Enlarged insets are shown (*bottom*). (D) Confocal images of immunostained CatSper1 (*top*) and CatSperζ (*bottom*) in 293T cells. FLAG-tagged CatSper1 (CS1-FLAG) and CatSperζ (CSζ-FLAG) were transiently expressed in 293T cells stably expressing WT (*middle*) and mutant (*right*) CatSperτ. Non-transduced 293T cells were used for negative control (*left*). Magnified insets are shown (*top right*). Hoechst to counterstain DNA (B, C, and D). (E) Quantitative analysis of CatSperτ-interactome from WT and *CatSpert*^Δ/Δ^ testis. Each identified protein represented as a dot was mapped according to its spectra-intensity in WT (y-axis) and *CatSpert*^Δ/Δ^ (x-axis). The fold change of CatSperτ (red dot) in WT over *CatSpert*^Δ/Δ^ testis was 2.74. The proteins with the fold changes above this value were considered to interact with CatSperτ significantly (blue dots, N=118) in testes. (F) Functional annotation of CatSperτ-interacting proteins. Enriched biological process gene ontologies are represented as bubbles and their functional categorization was visualized by a semantic similarity-based scatter plot. Bubble colors and sizes indicate false discovery rate (FDR) and proportion of total proteins annotated with the gene ontologies, respectively. (G) Proposed model for CatSperτ-mediated flagellar targeting of the CatSper channel. CatSperτ complexes with the vesicles carrying the channel complex, localizes them close to basal body and, and transports them to the elongating flagella of the developing spermatids in a quadrilateral formation. See also Figures S6 and S7, and Tables S1 and S2.

Our results highlight that CatSperτ has pivotal roles in intracellular transport and trafficking of the CatSper channel to the flagellar and nanodomains via the membrane-associating C2 domain (Figures 3, 5, 6, and S3, S5, S6). We hypothesized that CatSperτ mediates the interaction between the membrane vesicles and the motor proteins for translocalization of the cargoes. i.e., the CatSper channel. Thus, we identified CatSperτ interactome in testes by using immunoprecipitation (IP)-based mass spectrometry and performed functional annotation (Figures 7E, 7F, and S7E-S7I; Tables S1 and S2). A total 811 proteins were identified from the IP with α-CatSperτ-359 (WT, N=704; *CatSpert*^Δ/Δ^, N=706) and IP with normal IgG (WT, N=193) (Figure S7E; Table S1). 117 proteins with fold changes above the fold change of CatSperτ in WT over *CatSpert*^Δ/Δ^ testis (WT/*CatSpert*^Δ/Δ^ = 2.74) were considered to significantly associate with CatSperτ (Figures 7E and S7F). It is especially noted that we identified the membrane vesicle-associated proteins (ARF5, COPB1, COPB2, RAB11B, and RAB5B), motor proteins (MYO6, MYO7A, and DYNC1I2), and ciliary/flagellar proteins (TRAF3IP1, DYNC2LI1, and IFT140) from the CatSperτ interactome, suggesting CatSperτ manages transportation of cargoes in membrane vesicles. The pathway analysis of the CatSperτ interactome by Ingenuity Pathway Analysis further revealed that protein targeting and vesicle movement were enriched in CatSperτ interactome (Figure S7G). Functional annotation of gene ontologies showed that the biological process terms, such as protein targeting, vesicle-mediated transport, and intraciliary transport, were significantly enriched (Figures 7F and S7H; Table S2). All these results further support the predicted CatSperτ functions to modulate cargo transportation. In addition, the enriched molecular function and cellular component ontologies represent that the CatSperτ interactome proteins have binding ability to protein-containing complex or motor activity, and localize at ciliary basal body, the transition zone, or vesicle membrane (Figures S7H and S7I). We propose that CatSperτ is a molecular linker to connect CatSper-carrying vesicles and the motor proteins, thereby enabling the intracellular transportation, flagellar targeting, and quadrilinear arrangement of the CatSper channel complexes in developing spermatids (Figure 7G).

## Discussion

### CatSperτ Targets CatSper Channel to the Flagella by Adopting Ciliary Trafficking Machinery

Cilia and flagella are the specialized cellular projection that extend from the cell body via the shared structural core, the axoneme. Both cellular compartments are equipped with membrane receptors and ion channels, and function as signal receivers in eukaryotic systems (Nachury and Mick, 2019; Wachten et al., 2017). Our comparative mass spectrometry analysis of CatSperτ immunocomplex from WT and the C2-truncated CatSperτ testes identified proteins mainly involved in vesicle transportation and ciliary trafficking (Figure 7), suggesting CatSperτ adopts conventional ciliary trafficking machinery. Analogous to ARF4 and ARF-like ARL6 that recognize and package the cargoes for ciliary targeting (Deretic et al., 2005; Jin et al., 2010), CatSperτ interactome contains ARF5, RAB11B, and a RAB11 effector RAB11FIP1, which are likely to sort and package the CatSper complexes into carrier vesicles. Cytoplasmic dynein or myosin motors normally deliver cargo-carrying vesicles to the ciliary base directly from the Golgi or via pericentriolar endosomes (Morthorst et al., 2018). We also found cytoplasmic myosin (MYO6 and MYO7A) and dynein (DYNC1I2) motors in the CatSperτ interactome. We propose that these cytoplasmic motors further deliver CatSper-carrying vesicles to the flagellar base to dock and fuse cargoes to the periflagellar membrane. As previously shown for several GPCRs (i.e., SSTR3, MCHR1, and GPR161) which rely on IFT-A and TULP3 molecules to enter the cilia (Mukhopadhyay et al., 2010; Mukhopadhyay et al., 2013), we found an IFT-A component (IFT140) in the CatSperτ associated proteins, presumably to take on the CatSper complex from the periflagellar region to the flagellar membrane.

In developing spermatids, CatSperτ showed polarized distribution along the flagellum (Figure 5). Furthermore, CatSperτ was enriched at the pericentriolar region when expressed in heterologous systems (Figure 7). These localization patterns support that CatSperτ can mediate between CatSper cargoes and the early endosome to transport the channel into the flagellum. Together with CatSperτ-dependent localization of CatSper1, these results suggest that CatSperτ contributes to polarized transportation of cargo. CatSperτ, however, likely coordinates CatSper flagellar targeting together with additional players in the native system as small fractions of CatSper channels still traffic to the flagellum in *CatSpert*-mutant spermatids (Figure 5).

### CatSperτ is a Major CatSper Component in Organizing the Ca^2+^ Signaling Nanodomains

A functional CatSper channel can be assembled without either CatSperζ-EFCAB9 binary complex or CatSperτ, demonstrating that the contribution of these non-TM subunits to the channel assembly is more likely to be marginal (Chung et al., 2017; Hwang et al., 2019; see also Figure 4). For example, although discontinuous, four CatSper nanodomains still remain in *Efcab9*-null sperm that express CatSper channel containing CatSperτ, indicating that CatSperζ-EFCAB9 is dispensable for the channel trafficking to each quadrant in mature spermatozoa (Chung et al., 2017; Hwang et al., 2019). Compared to the absence of EFCAB9-CatSperζ, we note that CatSperτ loss-of-function reduces the protein levels of the CatSper complex more significantly and disarranges the channel localizations more severely in mature spermatozoa (Figures 1 and 3; see also Chung et al., 2017; Hwang et al., 2019). CatSperτ can localize in a quadrilinear fashion along the flagellum even without the assembled CatSper channel (Figure 6). The C2-truncated mutant prevents the CatSper channel from forming the four linear domains in the spermatids (Figure 6). Furthermore, CatSperτ is able to regulate CatSperζ intracellular localization in heterologous systems (Figure 7). Therefore, among the CatSper components reported so far, CatSperτ is the major component that targets the channel complex to flagella and organizes the quadrilinear nanodomains.

Genetic abrogation of the CatSper pore-forming or the auxiliary TM subunits results in entire loss of the CatSper channel complex in mature spermatozoa (Wang et al., 2021). This “all-in or all-out” expression pattern indicates that the pore-forming and auxiliary TM subunits are required to form a functional CatSper channel unit, also demonstrated by a recent atomic structure of the CatSper channel complex isolated from mouse sperm (Lin et al., 2021). The same study also reported CatSperτ (C2CD6) as one of the high confidence proteins from the mass spectrometry analysis of the purified CatSper complexes from mouse testes and epididymis. Yet, the resolution of the structures for the intracellular domains was low and did not resolve the intracellular components of the CatSper channel other than EFCAB9-CatSperζ (Lin et al., 2021). Whether the unassigned, intracellular electron densities from the existing structure include CatSperτ remains to be elucidated. Our recent studies also suggest an E3 ubiquitin-protein ligase, TRIM69, as another new CatSper component; its protein level and intraflagellar distribution in epididymal sperm is CatSper-dependent (Hwang et al., 2019; Zhao et al., 2021). Of note, our CatSperτ interactome from testes also includes TRIM69 (Table S1). Human CatSperτ is predicted to have ubiquitinated lysine (Akimov et al., 2018). Therefore, TRIM69 might transiently associate with CatSperτ, i.e., for post-translational modification, and contribute to the CatSper channel targeting and/or nanodomain organization.

### Membrane-Associating C2 Domain is Essential for CatSperτ Function

Diverse C2 domain containing proteins function in membrane trafficking, fusion, and signal transduction through its membrane-association and/or Ca^2+^ sensing ability (Brunger et al., 2018; Corbalan-Garcia and Gomez-Fernandez, 2014). Multiple ciliopathy genes also encode C2-domain containing proteins, highlighting the functional importance of C2 domain in modulating ciliary cargo transportation (Zhang and Aravind, 2012). For example, mutation of *MKS6* encoding CC2D2A causes Meckel syndrome. CC2D2A is localized at the transition zone, and its mutation prevents ciliary trafficking of the membrane proteins in mammalian cells (Garcia-Gonzalo et al., 2011). Another C2 protein, C2CD3, localizes at the centrosome and recruits ciliary proteins to dock vesicles to the mother centriole (Ye et al., 2014). C2CD3-interacting CEP120 contributes to form centriole appendages (Tsai et al., 2019) and its C2 domain mutation causes ciliopathy (Joseph et al., 2018).

C2 domain is composed of eight β-strands that form a conserved barrel structure (Nalefski and Falke, 1996; Nishizuka, 1988). The loops between β-strands form the membrane-associating region, which interacts with membranal phosphate via their positive charged or Ca^2+^-bound acidic residues (Corbalan-Garcia and Gomez-Fernandez, 2014). Our *CatSpert*-mutant mice clearly demonstrate that the C2 domain truncation, which is predicted to lose the first loop, is sufficient alone to impair the polarized localization of CatSperτ and its interaction with the vesicular components, resulting in the failure of the CatSper complex trafficking to the flagella (Figures 5 and 7). As most ciliary C2 domain proteins stay at the transition zone, it is intriguing that CatSperτ traffics not only to the flagella but also localizes into the four CatSper nanodomains. It is possible that the CatSperτ C2 domain also plays a role in Ca^2+^ sensing and/or domain stabilization in mature sperm flagella, which remains to be further investigated.

### The CatSper Levels Determine Sperm Capability to Maintain Ca^2+^ Homeostasis and Motility

Ca^2+^ is crucial for endurance and dynamic regulation of sperm motility. Ca^2+^ overload (Sanchez-Cardenas et al., 2018; Tateno et al., 2013) as well as depletion (Hwang et al., 2019; Marquez et al., 2007) renders sperm less motile. Under physiological conditions, Ca^2+^ homeostasis in mature spermatozoa would be mainly achieved via the balance between CatSper-mediated Ca^2+^ influx and PMCA4-mediated Ca^2+^ extrusion. [Ca^2+^]_i_ in *Pmca4*-null sperm is maintained abnormally high (Schuh et al., 2004). CatSper-deficient sperm gradually lose their motility under both non-capacitating and capacitating conditions (Hwang et al., 2019; Qi et al., 2007). In the current study, we find that there are critical levels of the CatSper proteins and *I_CatSper_* which can replenish [Ca^2+^]_i_ to recover the diminished motility when [Ca^2+^]_i_ is depleted. Sperm lacking EFCAB9-CatSperζ contain ∼30% of CatSper proteins and conduct around half of *I_CatSper_* compared to WT sperm, presumably due to the loss of gate inhibition (Chung et al., 2017; Hwang et al., 2019). These levels were sufficient to recover the motility of the mutant sperm when the available channels are stimulated. By contrast, *CatSpert*^Δ/Δ^ sperm retain ∼10% of CatSper proteins but conduct only ∼6% of *I_CatSper_* compared to WT sperm. Apparently, this level of CatSper activity was insufficient to fully recover sperm motility or swimming speeds, even under the capacitating condition that activates the channel (Figure 4). Human sperm from subfertile patients show significantly reduced Ca^2+^ influx and [Ca^2+^]_i_ after inducing CatSper activation (Kelly et al., 2018). Therefore, spermatozoa with more than this critical threshold of functional CatSper channels and the nanodomain structure are likely to manage the spatiotemporal Ca^2+^_i_ changes during their long travel to fertilize eggs in the female reproductive tract (Chung et al., 2014; Ded et al., 2020). Therefore, the levels of CatSper proteins and/or *I_CatSper_* can serve as a prognosis to determine clinical approaches, i.e., IVF vs. ICSI.

In summary, we have provided fundamental insights into the flagellar targeting mechanisms of the CatSper channel and its quadrilinear nanodomain organization (Figure 7G). We have identified CatSperτ as a new CatSper component and demonstrated that the C2-domain containing CatSperτ is a key regulator for CatSper flagellar targeting by mediating the channel complex and ciliary trafficking machineries. CatSperτ is in complex with the assembled CatSper channel and traffics the channel complex to developing spermatid flagella in a C2-dependent manner. CatSperτ interacts with vesicle and motor proteins, which are expected to transport CatSper-carrying vesicles to flagella in developing spermatids. We further show that CatSperτ is required for the quadrilinear arrangement of the targeted CatSper channel, which is crucial for sperm hyperactivation and male fertility. This study highlights the pivotal roles of CatSperτ in flagellar targeting and quadrilinear arrangement of the CatSper channel complex.

## Acknowledgement

We thank Jong-Nam Oh for assistance in flagellar waveform analysis and immunocytochemistry with human sperm samples, David E. Clapham and Byoung-Il Bae for sharing antibodies. This work was supported by start-up funds from Yale University School of Medicine, National Institute of Child Health and Human Development (R01HD096745), and Grantham Foundation to J.-J.C.; the Ministry of Education, Culture, Sports, Science, and Technology (MEXT)/Japan Society for the Promotion of Science (JSPS) KAKENHI grants (JP19H05750 and JP21H05033), and National Institute of Child Health and Human Development (R01HD088412 and P01HD087157) to M.I.; and the MEXT/JSPS KAKENHI grant (JP18K16735) to Y.L.; J.Y.H. is a recipient of postdoctoral fellowship from Male Contraceptive Initiative.

## Author contributions

J.-J.C. conceived and supervised the project. J.-J.C. and J.Y.H. designed, performed, and analyzed experiments. J.-J.C. and J.Y.H. generated *CatSpert* mutant mice. Y.L. and M.I. created *CatSpert* knockout mice. J.Y.H. performed molecular and cell biology experiments including expression construct generation, virus particle production, immunoprecipitation, immunocytochemistry, confocal and SIM imaging, sperm motility analysis, IP mass spectrometry and proteomics data analysis. H.W. did electrophysiological recordings, and H.W. and J.Y.H. analyzed the electrophysiology data. Y.L. performed IVF and histology experiments. M.I. provided crucial reagents and materials. J.Y.H. assembled figures. J.Y.H. and J.-J.C. wrote the manuscript with the input from the co-authors.

## STAR METHODS

### KEY RESOURCES TABLE

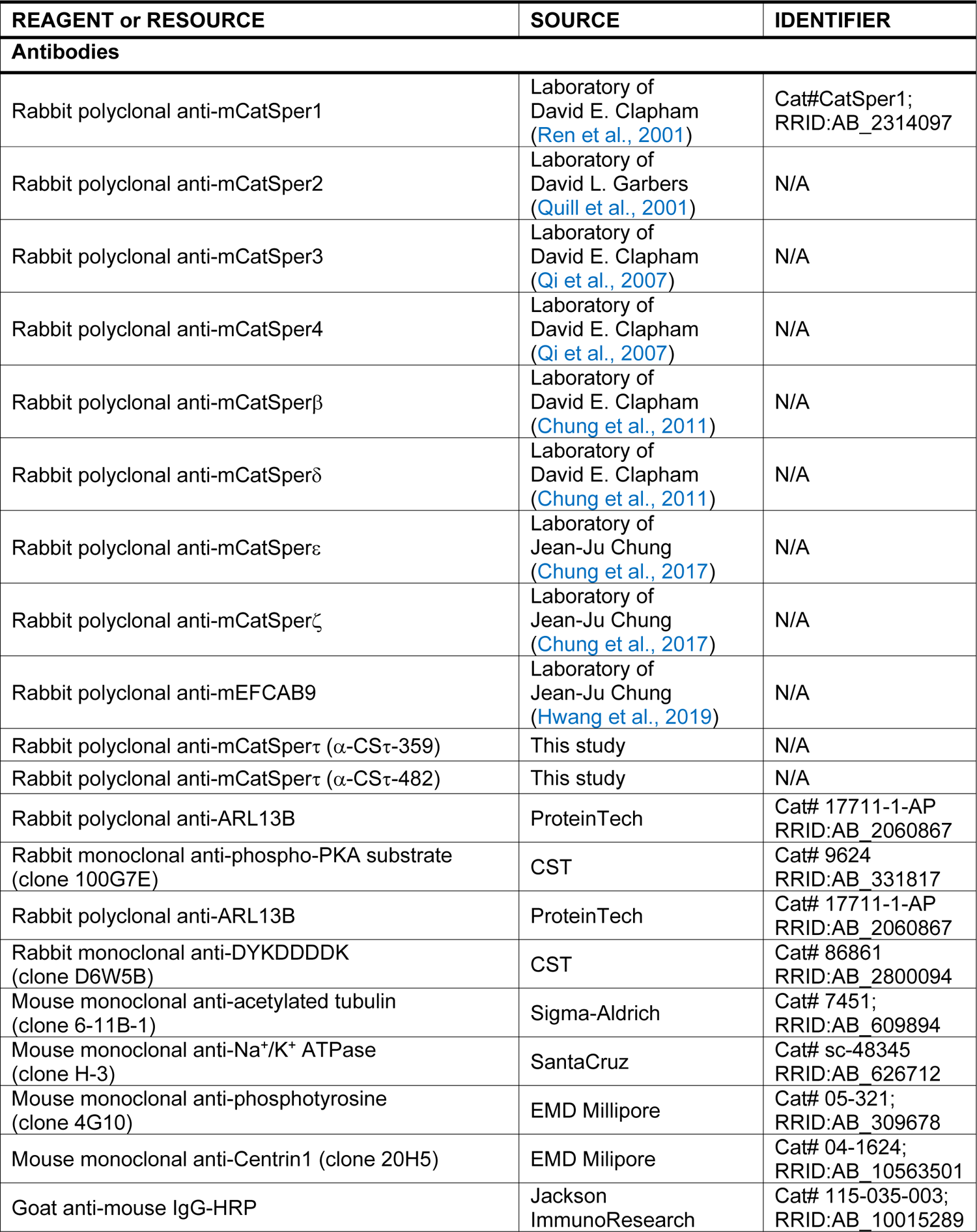
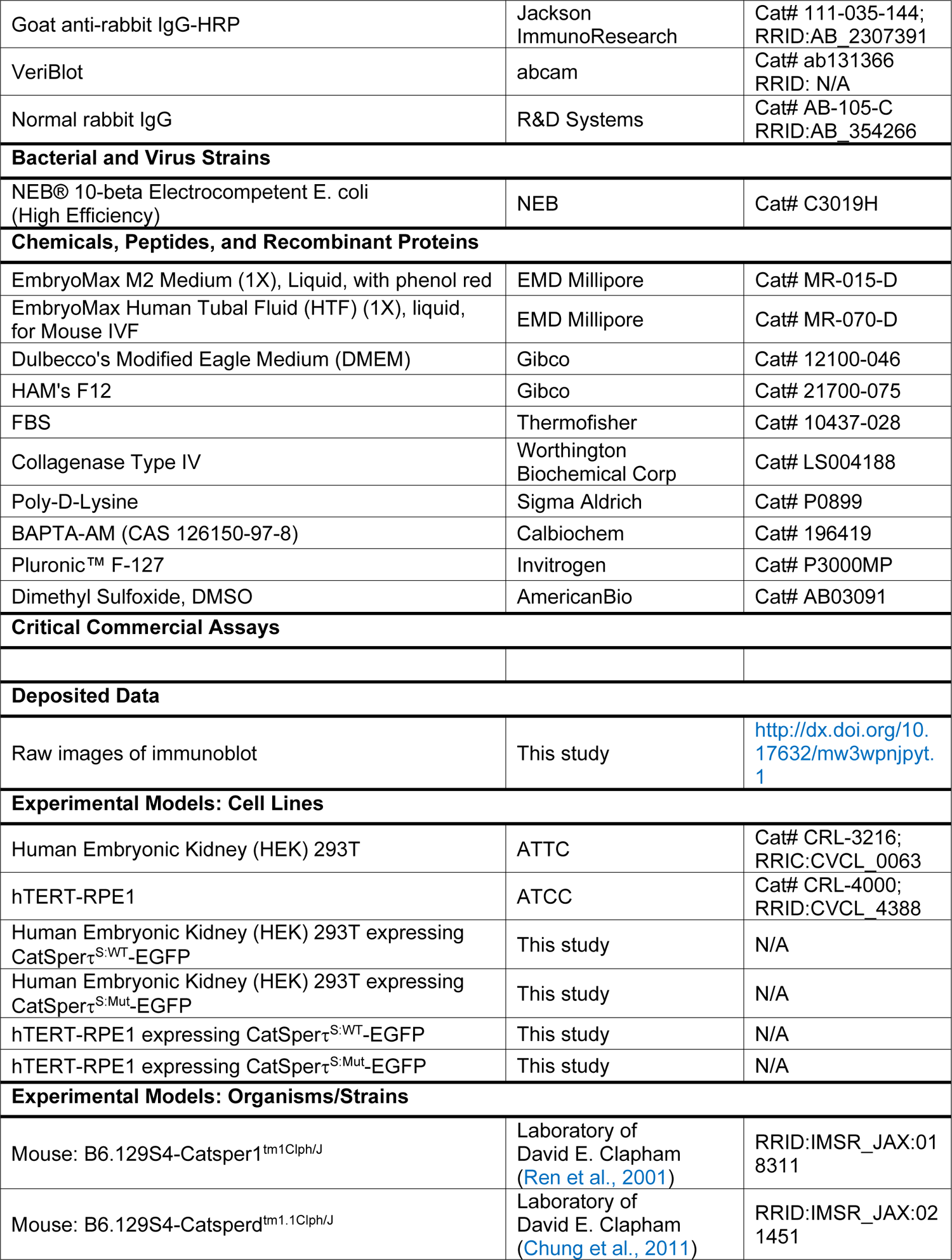
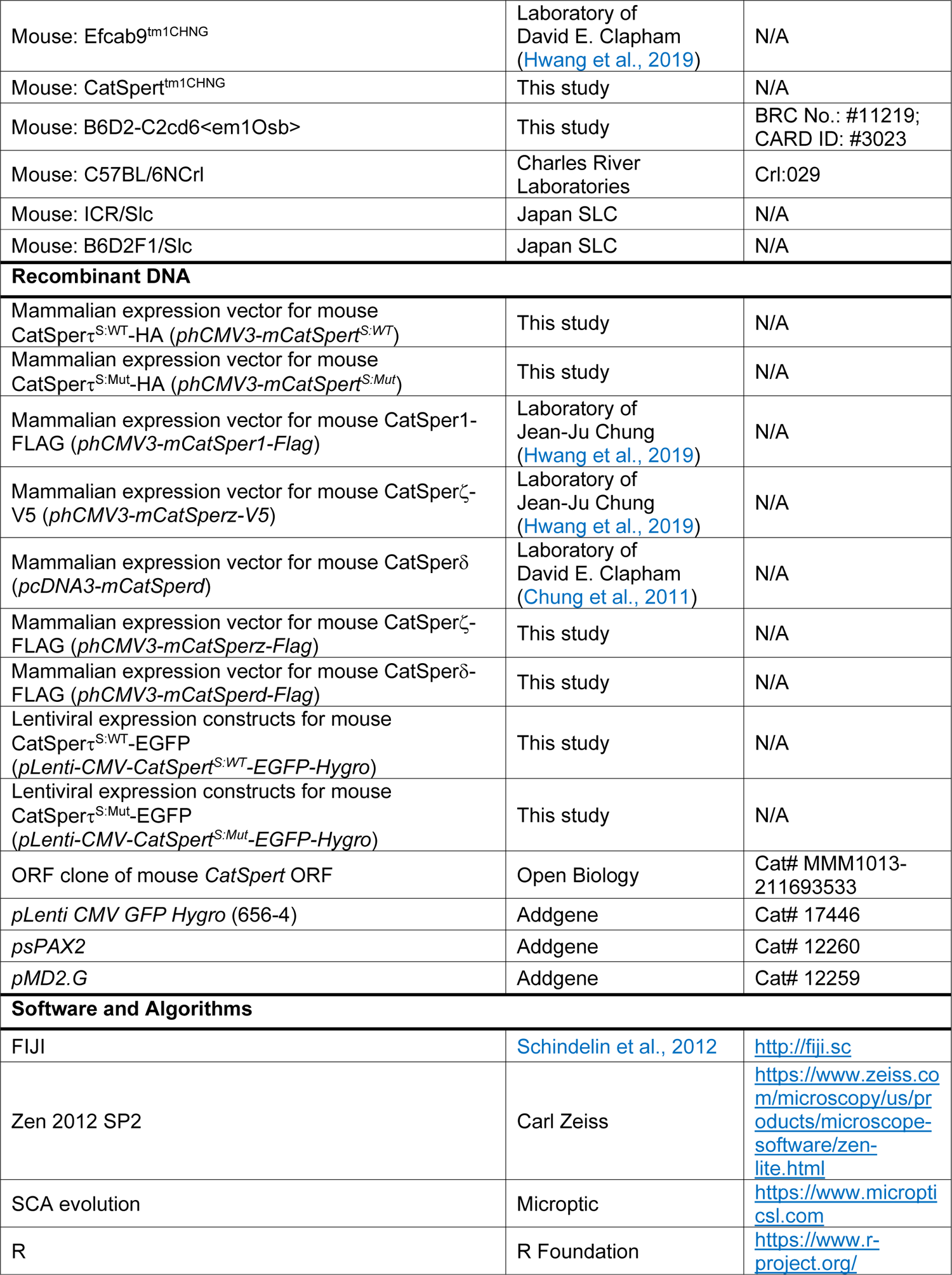
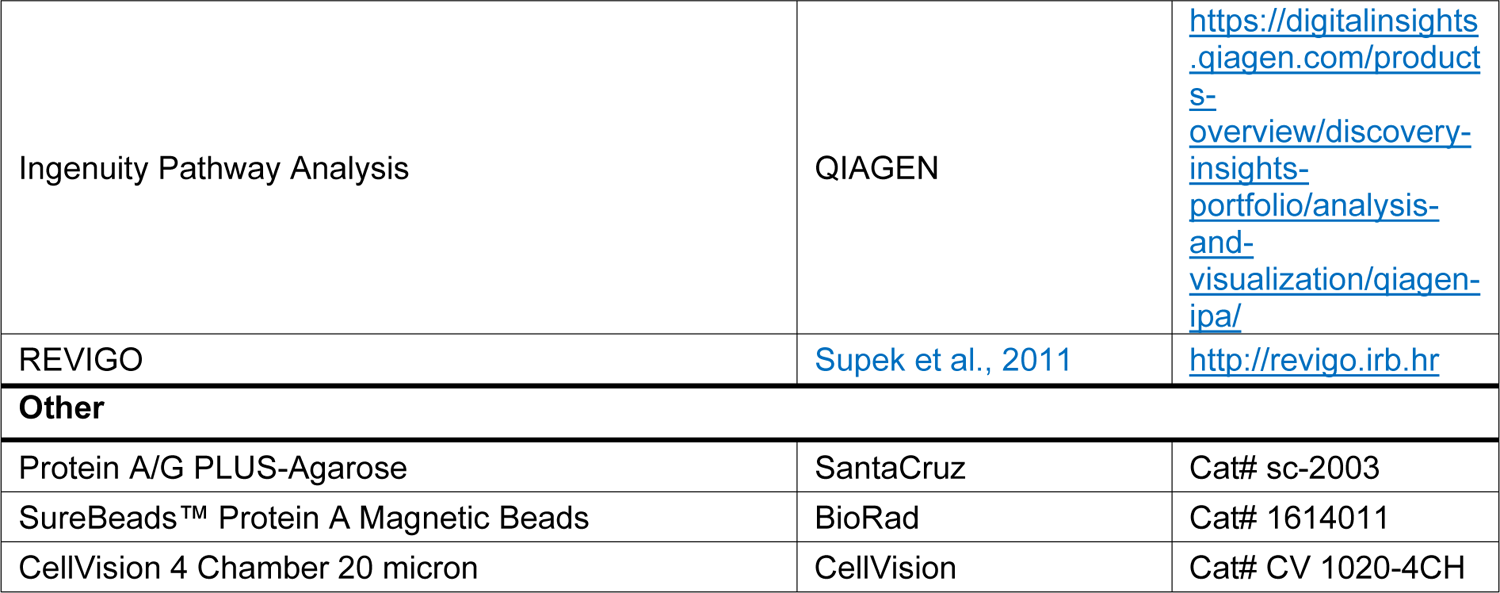

#### CONTACT FOR REAGENT AND RESOURCE SHARING

Further information and requests for the resources and reagents should be directed to the Lead Contact, Jean-Ju Chung.

### EXPERIMENTAL MODEL AND SUBJECT DETAILS

#### Animals

*CatSper1*, *CatSperd*, and *Efcab9*-null mice (Chung et al., 2011; Hwang et al., 2019; Ren et al., 2001) are maintained on a C57BL/6 background. WT C57BL/6 mice were purchased from Charles River Laboratory. WT B6D2F1 and ICR mice were purchased from Japan SLC. Mice were cared according to the guidelines approved by Institutional Animal Care and Use Committee (IACUC) for Yale University (#20079) and by the Animal Care and Use Committee of the Research Institute for Microbial Diseases, Osaka University (#Biken-AP-R03-01-0).

#### Generation of CatSpert-mutant and knockout mice by CRIPSR/Cas9 and Genotyping

*CatSpert*-mutant and knockout mice were generated on C57BL/6 and B6D2F1 background, respectively, using CRISPR/Cas9 method. For *CatSpert*-mutant mice, two guide RNAs (gRNAs), 5’-GCCACTCACGGACAAGAACA-3’ and 5’-TCTCCAAATGGTATCAAGTT-3’, in px330 were injected into pronuclei of the fertilized eggs obtained from super-ovulated females after mating with males. 2-cell embryos were transplanted into pseudo-pregnant females and founders’ tails were biopsied to extract genomic DNA (gDNA). To examine the large truncation at target region by CRSIPR/Cas9 editing, PCR was performed with F1 (5’-AAGACAGCTCCTGAGACGTG-3’) and R1 (5’-GGGGGTTGGAGTATGGAGGA-3’) primers. The PCR products were Sanger sequenced; founder females, which carry mutant allele with 128 or 134 bp deletion at coding region (*128del* and *134del*, respectively) were mated with WT C57BL/6 males to test germline transmission of the mutant allele. Two mutant mice lines were maintained, and genotyping was performed with F1-R1 primer pair for mutant allele and F2 (5’-ATCCACTGCCACTGCCTAGA-3’)-R1 primer pair for WT allele. For *CatSpert*-knockout mice, two CRISPR RNAs (crRNAs), 5’-AGCGCTCCACCTACGCCTAC-3’ and 5’-TCAGAGGTCTTCGGTAAATG-3’, were annealed to SygRNA^®^ Cas9 Synthetic trans-activating CRISPR RNA (tracrRNA; Sigma Aldrich), and then mixed with TrueCut™ Cas9 Protein v2 (Thermofisher) to form crRNA:tracrRNA:Cas9 ribonucleoprotein (RNP) complexes. The RNP complexes were introduced into the fertilized eggs obtained from super-ovulated WT B6D2F1 females by electroporation using an NEPA21 electroporator (Nepagene). Likewise, the treated eggs that developed into the 2-cell stage were transplanted into pseudo-pregnant ICR females, and gDNA was obtained from the founder animals by toe clipping. PCR was carried out using F1’ (5’-TGCACGTGGTCCAGGAAA-3’) and R1’ (5’-TTGCTGGGGGAGACTCACTTA-3’) primers or F2’ (5’-TGGTGCAGTATGGTGATAAGG-3’) and R1’ primers to detect the knockout or WT allele, respectively. The PCR products from the knockout allele were subjected to Sanger sequencing to verify the detailed deletion size.

### Cell Lines

#### Mammalian cell lines

HEK293T cells (ATCC; derived from female embryonic kidney) were cultured in DMEM (GIBCO) containing 10% FBS (Thermofisher) and 1x Pen/Strep (GIBCO) at 37 °C, 5% CO2 condition. hTERT-RPE1 (ATCC; derived from female retina pigmented epithelium) were cultured in 1:1 mixture of DMEM and Ham F12 (DMEM/F12, GIBCO) supplemented with 10% FBS (Thermofisher), 1x Pen/Strep (GIBCO), and 10 μg/ml Hygromycin B (Invitrogen) at 37 °C, 5% CO2 condition. HEK293T cells and hTERT-RPE1 cells stably expressing CatSperτ^S:WT^ and CatSperτ^S:Mut^ were cultured in the mediums for HEK293T cells and hTERT-RPE1 cells containing 50 μg/ml and 20 μg/ml Hygromycin B (Invitrogen), respectively.

#### Bacterial strains

NEB^®^ 10-β and NEB^®^ Stable (NEB) bacterial strains were used for molecular cloning.

### Sperm Preparation

#### Mouse sperm preparation and In Vitro Capacitation

Epididymal spermatozoa were collected from cauda, corpus, and cauda epididymis of adult male mice by swim-out methods in M2 medium (EMD Millipore) or HEPES-buffered saline (HS; Chung et al., 2017), or direct release into Toyoda-Yokoyama-Hosi (TYH) medium. Collected cauda sperm were incubated in human tubular fluid (HTF; EMD Millipore) at 2.0 – 3.0 x 10^6^ cells/ml concentration to induce capacitation for 90 minutes at 37 °C, 5% CO2 condition. For *in vitro* fertilization (IVF), sperm were capacitated in TYH medium at a concentration of 2.0 x 10^6^ cells/ml for 120 minutes at 37 °C, 5% CO2 condition to induce capacitation.

#### Human sperm preparation

Frozen human sperm vials from healthy donors were purchased from Fairfax Cryobank. The vials were thawed and mixed with warm HS. After washing with HS two times, human sperm were placed on top of 20% Percoll (Sigma Aldrich) in HS and incubated at 37 °C for 30 minutes. Immotile sperm in top layer were discarded and motile sperm were collected by centrifugation at 2,000 x g, followed by resuspension with HS.

### Testicular Germ Cell Preparation

#### Testicular cells preparation

Testes from adult male mice were collected and tunica albuginea, a capsule layer, was removed to harvest seminiferous tubules. Collected seminiferous tubules were washed with ice-cold PBS two times and chopped to dissociate germ cells. Dissociated testicular cells and chopped seminiferous tubules in PBS were filtered with cell strainer with 40 μm mesh size (Fisher Scientific). Filtered testicular cells were used for immunostaining.

#### STA-PUT germ cell separation

STA-PUT velocity sedimentation was applied to separate spermatogenic cell types (Bryant et al., 2013). Seminiferous tubules were collected from six to eight testes and washed with ice-cold PBS two times and DMEM (GIBCO) one time. Pulled seminiferous tubules were dissociated by incubation in 1 mg/ml collagenase Type IV (Worthington Biochemical Corp) at 37 °C for 10 minutes. Dissociated tubules were washed two times with DMEM by centrifugation at 500 x g for 5 minutes and incubated in EDTA-free 0.25% trypsin (GIBCO) with 15 unit/ml DNase 1 (NEB) at 37 °C for 10 minutes to dissociate testicular cells. Dissociated testicular cells in trypsin were mixed with DMEM (GIBCO) containing 10% FBS (Thermofisher) and 4 unit/ml DNase 1 (NEB), followed by filtering with 100 μm-pore cell strainer (Fisher Scientific). The filtrates were centrifuged and washed with DMEM (GIBCO) containing 10% FBS (Thermofisher) and 0.5% BSA (Sigma Aldrich) at 500 x g for 5 minutes. Washed germ cells were filtered with cell strainer with 40 μm mash pore. 2.0 – 3.0 x 10^8^ cells were loaded on top of 2 – 4% of BSA (Sigma Aldrich) gradient in DMEM (GIBCO) and sedimented for 2 hours to separate different types of testicular cells. Medium was fractioned and morphology of testicular cells in each fraction was examined using Axio observer Z1 microscope (Carl Zeiss) to enrich round spermatids.

## METHOD DETAILS

### Antibodies and Reagents

Rabbit polyclonal antibodies to recognize mouse CatSper1 (Ren et al., 2001), 2 (Quill et al., 2001), 3, 4 (Qi et al., 2007) β, δ (Chung et al., 2011), ε, ζ (Chung et al., 2017), and EFCAB9 (Hwang et al., 2019) were described previously. Polyclonal CatSperτ antibodies were generated by immunizing rabbits with KLH carrier-conjugated peptides corresponding to 359^th^ – 377^th^ (EKLREKPRERLERMKEEYK, α-τ-359; Open Biosystems) or 482^nd^ – 500^th^ (QIVEENEMPHLPKTSEPED, α-τ-482; Sigma Aldrich) residues of CatSperτ. Antisera from the immunized rabbits were affinity-purified using the peptides crosslinked to AminoLink^TM^ Coupling Resin (Pierce). All other commercial antibodies and reagents used in this study are listed in the key resource table.

### RNA Extraction, cDNA Synthesis, and RT-PCR

Total RNA was extracted from adult WT and homozygous *CatSpert-134del* mutant male testes using RNeasy Mini kit (QIAGEN). 500 ng of the extracted RNA was used for cDNA synthesis using iScript^TM^ cDNA Synthesis kit (Bio-Rad) according to manufacturer’s instruction. cDNAs were subjected to end-point PCR using OneTaq^®^ 2X Master Mix (NEB) or quantitative PCR (qPCR) using iTaq Universal SYBR Green Supermix (Bio-Rad). Primers, F3 (5’-CGGTTAGTGGCAGATAGGGC-3’), R3, (5’-TTGCTCTGCAGAGAACCTGG-3’), F4 (5’-TTTCCCACCCAGTCATCTAA-3’), R4 (5’-CTTCATCTCGCCAAACCTAA-3’), F5 (5’-CCTTGGCCAACAGCGTTTTA-3’), R5 (5’-TCTTTCCAACCTTTCCCGGG-3’), F6 (5’-AGCATTGGGGTTCCTGAAGC-3’), and R6 (5’-TCAACTCTCAGGAGCCCAAAC-3’), were used for end-point PCR and qF1 (5’-CAAGGGTAAAGGCACAGGAA-3’), qR1 (5’-TTTATGTGAATCGCCAGACAG-3’), qF2 (5’-GACAAAAAGGGATAATAAGGGAAG-3’), and qR2 (5’-TGAAATAGCTTCATATTTTCTGTGATG-3’) were used for qPCR. *TBP* was used as a reference gene to normalize transcript levels by ddCt method.

### Molecular Cloning and Lentivirus Production

#### Transient expression constructs

Mammalian expression constructs for CatSper1 (*phCMV3-CatSper1*-*Flag*) was described in previous study (Hwang et al., 2019). Mouse *CatSperd* or *z* ORFs (Chung et al., 2011; Hwang et al., 2019) were subcloned into phCMV3 to express C-terminal FLAG-tagged CatSper subunits (*phCMV3*-*CatSperd* or *z-Flag*) using NEBuilder^®^ HiFi DNA Assembly Kit (NEB). A stop codon was placed at the upstream of HA-encoding sequences of phCMV3 vector for FLAG-tagged CatSper subunit cloning.

#### Lentiviral Transfer constructs

An ORF clone of mouse *CatSpert* (NM_175200) Open Biosystems, clone 40090362) was subcloned into phCMV3 (*phCMV3-CatSpert^S:WT^*) encoding short form of CatSperτ (CatSperτ^S:WT^) tagged with HA at C-terminus. A CDS of the mouse mutant CatSperτ (CatSperτ^S:Mut^) with C2 domain truncation (Δ103-163) was cloned into *phCMV3* vector by assembling two CDS fragments amplified from *phCMV3-CatSpert^S:WT^* (*phCMV3-CatSpert^S:Mut^*) using NEBuilder^®^ HiFi DNA Assembly Kit (NEB). ORF sequences of CatSperτ^S:WT^ (*phCMV3-CatSpert^S:WT^*) and CatSperτ^S:Mut^ (*phCMV3-CatSpert^S:Mut^*) were amplified using Q5^®^ Hot Start High-Fidelity 2X Master Mix (NEB) and subcloned into *pLenti-CMV-GFP-Hygro* (656-4) (gifted from Eric Campeau & Paul Kaufman; Addgene plasmid # 17446) using NEBuilder^®^ HiFi DNA Assembly Kit (NEB) (*pLenti-CMV-CatSpert^S:WT^-EGFP-Hygro* and *pLenti-CMV-CatSpert^S:Mut^-EGFP-Hygro*, respectively).

#### Lentivirus Production

Cloned lentiviral expression vectors (*pLenti-CMV-CatSpert^S:WT^-EGFP-Hygro* and *pLenti-CMV-CatSpert^S:Mut^-EGFP-Hygro*), and lentiviral packaging (*psPAX2*, Addgene plasmid # 12260) and envelop (*pMD2.G*, Addgene plasmid # 12259) plasmids (gift from Didier Trono) were transfected into cultured HEK293T cells using polyethylenimine (PEI). Culture medium was harvested after 1-3 days transfection. Collected medium with virus particles were centrifuged at 1,000 x g for 15 minutes at 4 °C and the supernatant were mixed with 4X lentivirus concentration solution −40% (w/v) polyethylene glycol 8000 (PEG 8000, Fisher Scientific), 1.2 M NaCl, 2.7 mM KCl, 10 mM Na2HPO4, and 1.8 mM KH2PO4, pH7.2 - followed by incubation at 4 °C for overnight. Precipitated virus particles were pelleted by centrifugation at 1,600 x g for 1 hour at 4 °C and resuspended with DMEM (GIBCO). Concentrated virus particles were frozen and stored at − 80 °C until use.

### Recombinant Protein Expression in Mammalian Cells

#### Transient protein expression

Mammalian expression vectors encoding FLAG-tagged mouse CatSper proteins (1, δ, and ζ) were transiently expressed in HEK293T cells. Cultured HEK293T cells were transfected with plasmid to express recombinant proteins using PEI. After 36 – 48 hours from the transfection, cells were used for immunostaining.

#### Stable protein expression

HEK293T and hTERT-RPE1 cells were transduced with the lentiviral particles to express EGFP-tagged mouse WT and mutant CatSperτ (CatSperτ^S:WT^-EGFP and CatSperτ^S:Mut^-EGFP, respectively). Briefly, HEK293T and hTERT-RPE1 cells were placed in DMEM containing 10% FBS, 10 μg/ml polybrene (EMD Millipore), and lentiviral particles, and subjected to cytospin centrifugation at 1,000 x g for 1 hour at room temperature (RT). The cells were incubated at 37°C, 5% CO2 for one day and the medium was changed to DMDM (HEK293T) or DMEM/F12 (hTERT-RPE1) containing 10% FBS and 1X Pen/Strep. The lentivirus-transduced HEK293T and hTERT-RPE1 cells to express mouse CatSpert^S:WT^-EGFP and CatSperτ^S:Mut^-EGFP were passaged one time and cultured in the FBS- and Pen/Strep-containing media supplemented with and 50 μg/ml (HEK293T cells) or 20 μg/ml (hTERT-RPE1) hygromycin B (Invitrogen). Stable protein expression was checked by epifluorescence microscope (ZOE Fluorescent Cell Imager, BioRad).

### Protein Solubilization, Extraction, and Immunoblotting

#### Testis microsome

Mouse testes were homogenized in 0.32M sucrose and the homogenates were centrifuged at 4 °C, 1,000 x g for 10 minutes to remove cell debris and nucleus. Supernatant were collected and centrifuged at 100,000 rpm for 60 minutes at 4°C to separate cytosolic (cyto) and microsome (mic) fractions in supernatant and pellet, respectively. Microsome proteins were solubilized in PBS containing 1% Triton X-100 and cOmplete Mini, EDTA-free Protease Inhibitor Cocktail (Roche) for 2 hours at 4°C with gentle rocking. The microsome lysates were centrifuged at 18,000 x g for 1 hour at 4 °C to obtain soluble (supernatant) and insoluble (pellet) fractions. The insoluble microsome fractions were further lysed with 2X LDS sampling buffer by vortexing for 10 minutes at RT followed by centrifugation at 18,000 x g for 30 minutes at 4 °C. Cytosolic and solubilized microsome proteins mixed to 1X LDS buffer and insoluble microsome proteins in 2X LDS buffer were volume-equivalented and denatured by boiling at 75°C with 50 μM at dithiothreitol (DTT) for SDS-PAGE and immunoblotting to examine the protein partitioning. Primary antibodies used for the immunoblotting were: rabbit polyclonal anti-mouse CatSperτ (α-CSτ-359 and α-CSτ-482, 1 μg/ml each) and mouse monoclonal anti-Na^+^/K^+^ ATPase (1:500; SantaCruz) and anti-acetylated tubulin (1:2,000; clone 6-11B-1, Sigma Aldrich). Goat anti-mouse or rabbit IgG conjugated with HRP were used for secondary antibodies (1:10,000; Jackson ImmunoResearch). Solubilized microsome fractions were also used for coIP.

#### Testicular germ cells

Round spermatids enriched by STA-PUT were lysed with 1% Triton X-100 in PBS supplemented with EDTA-free protease inhibitor cocktail (Roche) by gentle rocking at 4°C for 2 hours. The lysates were centrifuged at 18,000 x g for 30 minutes at 4°C and solubilized proteins in supernatant were collected. Solubilized proteins were subjected to coIP.

#### Epdidymal sperm cells

Whole sperm proteins were extracted as previously described (Hwang et al., 2019). Collected epididymal sperm from corpus and cauda were washed with PBS and lysed with 2X LDS by vortexing for 10 minutes at RT. The lysates were centrifuged at 4°C, 18,000 x g for 10 minutes. The supernatants were mixed to 50 μM DTT and denatured by boiling at 75°C for 10 minutes. Denatured sperm proteins were subjected to SDS-PAGE and immunoblotted. The used primary antibodies were: Rabbit polyclonal anti-mouse CatSper1, CatSper2, CatSper3, CatSper4, CatSperβ, CatSperδ, CatSperτ, and EFCAB9 at 1 μg/ml, mouse CatSperζ (2.7 μg/ml), and rabbit monoclonal anti-phospho-PKA substrate (clone 100G7E, CST, 1:1,000) and mouse monoclonal anti-phosphotyrosine (1 μg/ml; clone 4G10, Sigma Aldrich) and acetylated tubulin (1:20,000; clone 6-11B-1, Sigma Aldrich). HRP-conjugated goat anti-mouse IgG and goat anti-rabbit IgG were used for secondary antibodies (1:10,000; Jackson ImmunoResearch).

### Co-immunoprecipitation

Solubilized testis microsome and round spermatids were subjected to coIP. Solubilized proteins in lysis buffer were mixed with SureBeads^TM^ Protein A Magnetic Beads (BioRad) conjugated with either 1 μg of rabbit polyclonal anti-CatSper1, CatSperδ, or CatSperτ (α-CSτ-482), or 0.5 μl of rabbit monoclonal anti-DYKDDDDK (clone D6W5B, CST). After incubation for overnight at 4°C, the resins were washed with lysis buffer and eluted immunocomplexes were eluted with 2X LDS buffer supplemented with 50 μM DTT by boiling at 75°C for 10 minutes. The elutes were subjected to SDS-PAGE and immunoblotting. Primary antibodies for immunoblotting were: rabbit polyclonal anti-mouse CatSper1, CatSper2, CatSper3, CatSperβ, CatSperδ, CatSperε, and CatSperτ at 1 μg/ml, and rabbit monoclonal anti-DYKDDDDK (1:2,000; clone D6W5B, CST). For secondary antibodies, VeriBlot (1:200-1:500; Abcam) and HRP-conjugated goat anti-mouse or rabbit IgG (1:10,000; Jackson ImmunoResearch) were used.

### IP Mass Spectrometry and Proteomics Analyses

#### Sample preparation

Testis microsome of WT and *CatSpert*^Δ/Δ^ males were prepared and solubilized as described above. Lysates were centrifuged at 18,000 x g for 1 hour at 4°C and supernatants were mixed with Protein A/G PLUS-Agarose (SantaCruz) and incubated for 1 hour at RT to remove mouse immunoglobulin. Flow through were collected followed by incubation with anti-CatSperτ (α-CSτ−359; WT and *CatSpert*^Δ/Δ^) or normal rabbit IgG (R&D SYSTEMS; WT) crosslinked to Protein A/G PLUS-Agarose (SantaCruz) using 20 mM dimethyl pimelimidate (Sigma Aldrich) at 4°C for overnight. The resins were washed with 1% Triton X-100 in PBS and eluted two times by incubation with 0.1M glycine, pH2.3 at RT for 5 minutes. Elutes from five repeats were pulled and incubated with 4-volumes of acetone at – 20°C for overnight to precipitate proteins. The mixture was centrifuged at 18,000 x g for 1 hour at 4°C followed elution with 8M urea, pH7.4. The elutes in 1X LDS with 50 mM DTT were boiled at 75°C for 5 minutes and subjected to SDS-PAGE. The gels were run shortly and stained using Imperial™ Protein Stain (Thermofisher). Stained bands were cut subjected to LC-MS/MS

#### Protein LC-MS/MS

The gel pieces were washed and dehydrated with acetonitrile for 10 minutes. The gel pieces were dried and subjected to trypsin-digestion for overnight at 37°C. Digested peptides were reverse-phase eluted (Peng and Gygi, 2001) and subjected to mass spectrometric analysis using an LTQ Orbitrap Velos Pro ion-trap mass spectrometer (Thermofisher). Peptides were detected, isolated, and fragmented to produce a tandem mass spectrum of each peptide, which were matched with protein database using Sequest (Thermofisher) to identify peptide sequences. Data was filtered to between a one and two percent peptide false discovery rate. Identified CatSperτ-interactomes were subjected to pathway analysis and functional annotation by using Ingenuity Pathway Analysis (QIAGEN) and STRING Version 11 (https://string-db.org; Szklarczyk et al., 2019).

Enriched gene ontologies were functionally categorized by using REVIGO (http://revigo.irb.hr; Supek et al., 2011) and visualized by using R software.

#### Immunocytochemistry

Collected epididymal sperm, testicular cells, and cultured cells were subjected to immunocytochemistry. Epididymal sperm were washed with PBS and attached to glass coverslip by centrifugation at 700 x g for 5 minutes. The attached sperm cells were fixed with 4% PFA in PBS for 10 minutes at RT and washed with PBS three times. Dissociated testicular cells in PBS were fixed with PFA in 4% concentration for 5 minutes followed by centrifugation at 250 x g for 3 minutes to attach to coverslips coated with poly-D-lysine (Sigma Aldrich). Testicular cells were air-dried shortly. Cultured cells on glass coverslip coated with poly-D-lysine were washed with PBS and fixed with 4% PFA in PBS for 10 minutes at RT. The fixed epididymal sperm, testicular cells, and cultured cells on glass coverslips were permeabilized with 0.1% Triton X-100 in PBS for 10 minutes at RT and blocked with 10% normal goat serum in PBS for 1 hour at RT. Blocked samples were incubated with primary antibodies at 4°C for overnight. Primary antibodies used for the immunocytochemistry were: rabbit polyclonal anti-CatSper1 (10 μg/ml), CatSperδ (10 μg/ml), CatSperτ (α-CSτ-359, 20 μg/ml; α-CSτ-482, 10 μg/ml), and ARL13B (1:200, Proteintech), rabbit monoclonal anti-DYKDDDDK (1:200; clone D6W5B, CST), and mouse monoclonal anti-pTyr (1:100, clone 4G10, Sigma Aldrich), anti-acetylated tubulin (1:200, clone 6-11B-1, Sigma Aldrich), and anti-CENTRIN1 (1:100, clone 20H5, Sigma Aldrich). Immunostained samples were washed with 0.1% Triton X-100 in PBS one time and PBS two times followed by incubation with goat anti-rabbit or mouse IgG conjugated with Alexa 488 or Alexa 568 (1:1,000, Invitrogen) in blocking solution for 1 hour. After incubation with secondary antibodies, coverslips were mounted with Vectashield (Vector Laboratory) and imaged with Zeiss LSM710 Elyra P1 using Plan-Apochrombat 63X/1.40 and alpha Plan-APO 100X/1.46 oil objective lens (Carl Zeiss). Hoechst (Invitrogen) were used for counter staining.

### Structured Illumination Microscopy

Immunostained mouse epididymal sperm and elongated spermatids (step 13-16) in dissociated testicular cells were prepared as described in *Immunocytochemistry*. 3D structural illumination microscopy (SIM) imaging was performed with Zeiss LSM710 Elyra P1 using alpha Plan-APO 100X/1.46 oil objective lens (Carl Zeiss). Z-stack images was acquired with 100 or 200 nm intervals and each section was taken using images were taken using 5 grid rotations with a 51 nm SIM grating period and a laser at 561 nm wavelength. Raw images were processed and rendered using Zen 2012 SP2 software (Carl Zeiss).

### Mating Test and *In Vitro* Fertilization

Adult female mice with normal fertility were housed with heterozygous or homozygous *CatSpert* mutant or knockout males over two months, and the pregnancies and litter sizes were recorded.

For IVF, Cauda epididymal spermatozoa from WT or knockout males were released into the TYH medium drops at a concentration of 2.0 x 10^5^ cells/ml. After 2 hours of incubation at 37°C, 5% CO2 condition, the capacitated spermatozoa were transferred to new TYH medium drops containing cumulus-oocyte complexes (COCs) obtained from super-ovulated B6D2F1 females at a final sperm concentration of 2.0 x 10^5^ cells/ml. After 6 hours of incubation, the COCs were treated with hyaluronidase and washed in clean TYH drops repeatedly to denude the cumulus cells. To determine the fertilization success, abnormal or parthenogenic eggs were excluded and the ones bearing two pronuclei were counted as fertilized eggs.

### Histology Analysis

Testes from WT and *CatSpert*-knockout males were fixed in Bouin’s fluid (Polysciences), dehydrated in ethanol, embedded in paraffin wax, and sectioned at a thickness of 5 μm on an HM325 microtome (Microm). The paraffin sections were adhered to microscope slides, rehydrated with graded concentrations of ethanol, stained with 1% periodic acid (Nacalai) and Schiff’s reagent (Wako), and counter-stained with Mayer’s hematoxylin solution (Wako). After dehydration with graded concentrations of ethanol and xylene, the microscope slides were mounted with Entellan® new (Sigma Aldrich) mounting medium and observed under an Olympus BX-53 microscope.

### Sperm Motility Analysis

#### Flagellar Waveform Analysis

Sperm flagellar movement were analyzed as previous studies (Hwang et al., 2019; Hwang et al., 2021). In brief, non-capacitated or capacitated cauda epididymal sperm (2 x 10^5^ cells/ml) were placed into fibronectin-coated imaging chamber for Delta-T culture dish controller (Bioptech) filled with 37°C HEPES-buffered HTF medium (Chung et al., 2017). Flagellar movements of the sperm cells of which heads were tethered on the plate were recorded for 2 seconds with 200 frame per seconds (fps) speed using pco.edge sCMOS camera equipped in Axio observer Z1 microscope (Carl Zeiss). Original image stacks were rendered to movies. Beating frequency and α-angle of sperm tail were measured and overlaid-images of sperm flagellar waveform were generated by using FIJI software (Schindelin et al., 2012).

#### Computer-Assisted Sperm Analysis

Non-capacitated and capacitated sperm cells (3.0 x 10^6^ cells/ml) were loaded in 20 μm-depth of slide chamber (CellVision) and sperm motility was examined on 37°C warm-stage. Motile sperm were imaged with Nikon E200 microscope under 10x phase contrast objective (CFI Plan Achro 10X/0.25 Ph1 BM, Nikon) and recorded at 50 fps speed using CMOS video camera (Basler acA1300-200μm, Basler AG). The recorded movies were analyzed by Sperm Class Analyzer software (Microptic). Over 200 total sperm were imaged for each group.

#### Ca^2+^ handling Assay

Intracellular Ca^2+^-dependent sperm motility changes (Ca^2+^ handling assay) were examined as performed previous study (Hwang et al., 2019). Briefly, epididymal sperm from WT, *CatSperd*-null and *CatSpert*^Δ/Δ^ sperm in M2 medium (3.0 x 10^6^ cells) were loaded with 5 μM BAPTA-AM (bis-(o-aminophenoxy)ethane-N,N,N’,N’-tetra-acetic acid acetoxymethyl ester, Calbiochem) or vehicle (0.05% DMSO and 0.01% F-127) and incubated for 90 minutes at 37 °C. After incubation, sperm cells were washed with HS medium by centrifugation two times at 700 x g for 2 minutes to remove BAPTA-AM and eluted with HTF medium (EMD Millipore). Eluted sperm in HTF was incubated at 37°C, 5% CO2 condition for 90 minutes to restore intracellular Ca^2+^ level via activated CatSper channel by inducing capacitation. Motilities of over 200 sperm cells were imaged and analyzed at each time point as described in *Computer-Assisted Sperm Analysis*.

### Electrophysiology

Corpus epididymal sperm were washed and resuspended in HS medium followed by attached on the 35mm culture dish. Gigaohm seals were formed at the cytoplasmic droplet of sperm (Kirichok et al., 2006). Cells were broken in by applying voltage pulses (450-600 mV, 5 ms) and simultaneous suction. Whole-cell CatSper currents were recorded from WT, *CatSperd*^-/-^, and *CatSpert*^Δ/Δ^ sperm in divalent-free bath solution (DVF, 150 mM NaCl, 20 mM HEPES, and 5 EDTA, pH 7.4). Intrapipette solution for CatSper current recording consists of 135 mM CsMes, 10 mM HEPES, 10 mM EGTA, and 5 mM CsCl adjusted to pH 7.2 with CsOH. To induce intracellular alkalinization, 10 mM NH_4_Cl was added to bath solution by perfusion system. Data were sampled at 10 Hz and filtered at 1 kHz. The current data were analyzed with Clampfit (Axon, Gilze, Netherlands), and figures were plotted with Grapher 8 software (Golden Software, Inc., Golden, Colorado).

## QUANTIFICATION AND STATISTICAL ANALYSIS

Statistical analyses were carried out using Student’s t-test. Differences were considered significant at *p<0.05, **p<0.01, and ***p<0.001.

## DATA AND SOFTWARE AVAILABILITY

The raw images of immunoblotting in this study are deposited in Mendeley Data at http://dx.doi.org/10.17632/mw3wpnjpyt.1. All software used in this study is listed in the Key Resource Table.

## SUPPLEMENTAL INFORMATION

Supplemental Information includes seven figures, two tables, and a movie can be found with this article online at xxx.

### Supplementary Figures and Legends

**Figure S1.**
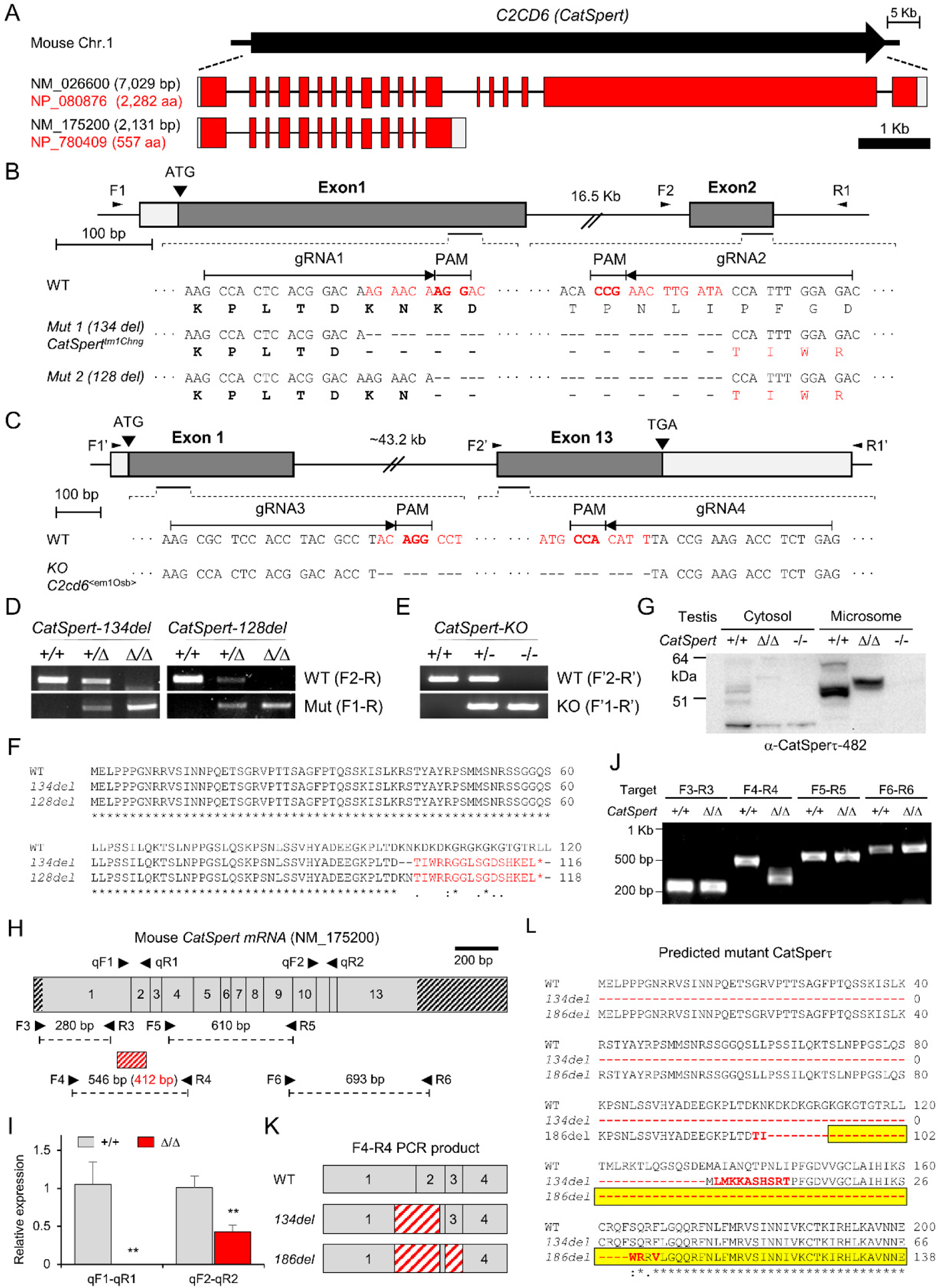
Generation of *CatSpert* deletion mutant and knockout mice, related to Figures 1 and 2. (A) Genome structure of mouse *C2CD6* gene encoding CatSperτ. Represented are two variants (NM_026600 and NM_175200) that encode the long (CatSperτ^L^, NP_080876) and short (CatSperτ^S^, NP_780409) isoforms, respectively. Marked in red (coding region) and gray (UTR) boxes are the exons in scale. (B-C) Generation of *CatSpert* deletion mutant (Δ/Δ) and knockout (-/-) mice using CRISPR/Cas9 genome editing. (B) Mutant alleles with 134 bp (*CatSpert*-*134del*) and 128 bp (*CatSpert-128del*) deletion in the CatSperτ coding region. The deletions are expected to cause out-of-frame mutations, resulting in early termination of protein translation from the start codon. (C) Knockout allele with 43 kb deletion at genomic region encoding CatSperτ. (D-E) Genomic DNA PCR for genotyping of *CatSpert*-mutant (D) and knockout (E) mice with the primer pairs as marked in panels B and C. (F) Alignment of the predicted amino acid sequences translated from WT and the two mutant alleles in B. Amino acids in red indicate non-native sequences from the frameshift before termination. (G) Detection of truncated CatSperτ protein in *CatSpert-134del*^Δ/Δ^ testes. Testis homogenates from WT, *CatSpert-134del*^Δ/Δ^, and *CatSpert* knockout mice were separated into cytoplasm (Cyt) and microsome (Mic) fractions and subjected to immunoblotting. (H-J) Detection of truncated *CatSpert* mRNA in *CatSpert-134del*^Δ/Δ^ testes. (H) Exon composition of the *CatSpert* transcript (NM_175200). Expected PCR product sizes for each primer pair are marked. A red box indicates the deleted coding region in *CatSpert-134del*^Δ/Δ^ mouse genome. (I) Real-time RT-PCR and (J) end-point RT-PCR analyses of *CatSpert* mRNA expression in *CatSpert*-*134del*^Δ/Δ^ testes. (K) Diagrams of the truncated *CatSpert* transcripts. Sanger sequencing of the F4-R4 PCR product reveals 134-bp (*middle*) or 186-bp (*bottom*) deleted *CatSpert* transcripts in *CatSpert-134del*^Δ/Δ^ testes. (K) Sequence alignment of the WT and predicted mutant CatSperτ sequences translated from the open reading frames of the truncated *CatSpert* transcripts. Red characters are the non-native amino acid sequences by deletion and frameshift. The sequence corresponding to C2 domain is yellow-boxed.

**Figure S2.**
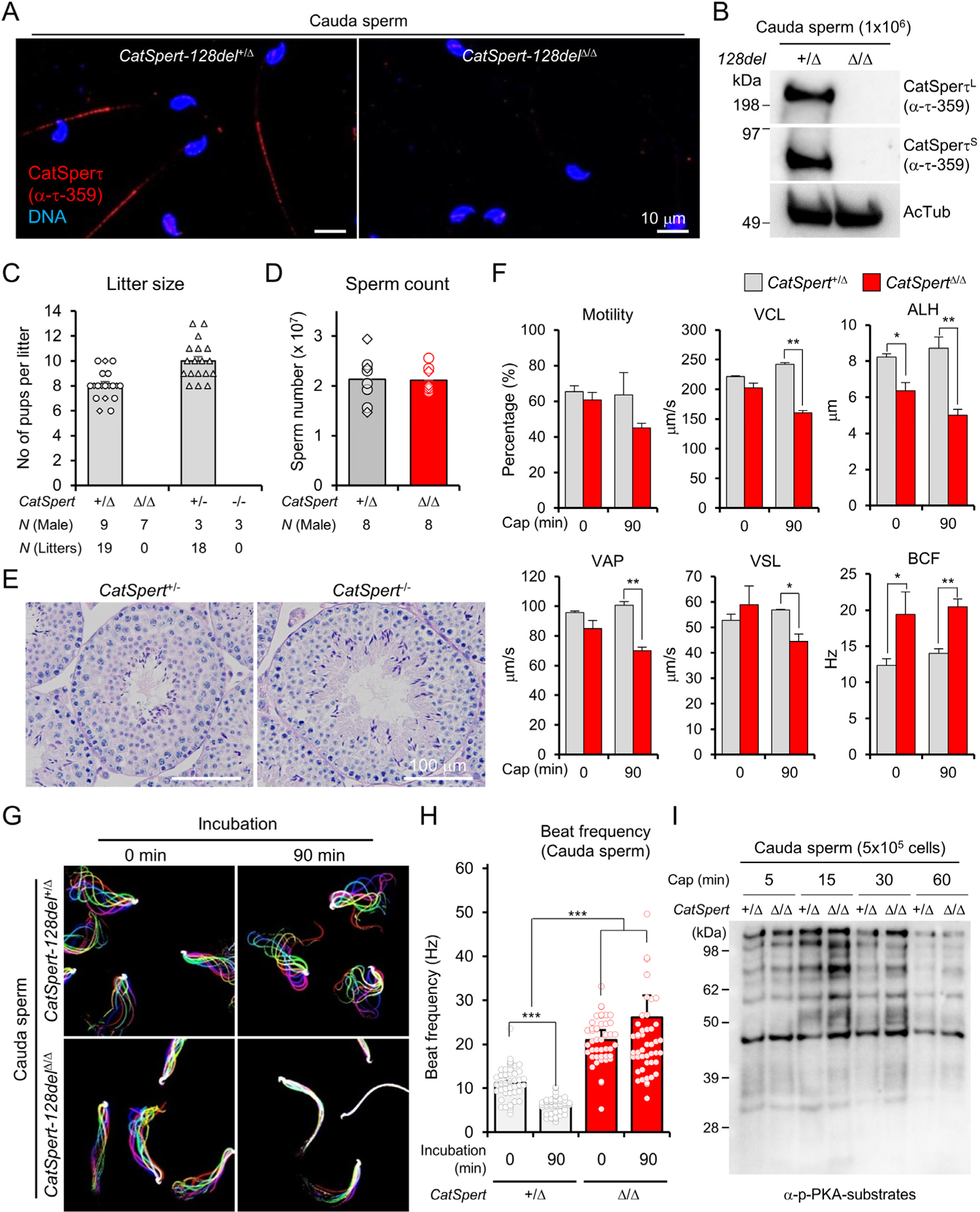
Characterization of *CatSpert* mutant and knockout mice, related to Figure 2. (**A-B**) Absence of the truncated CatSperτ in *CatSpert-128del*^Δ/Δ^ sperm by immunostaining (A) and immunoblotting (B). As *CatSpert-134del*^Δ/Δ^ and *CatSpert-128del*^Δ/Δ^ show identical phenotypes from initial characterization, *CatSpert-134del*^Δ/Δ^ mice were used as *CatSpert*^Δ/Δ^ throughout this study unless indicated. (C) Litter sizes from WT, heterozygous, and homozygous females mated with heterozygous and homozygous *CatSpert-*mutant (+/Δ, 8.1 ± 0.3; Δ/Δ, 0.0 ± 0.0) and knockout (+/-, 10.0 ± 0.4; -/-, 0.0 ± 0.0) males. Females are all fertile as shown. Litters from the females mated with *CatSpert-134del* (circles; +/Δ, N=6, litter number=12; Δ/Δ, N=4, litter number=0) and *128del* (triangles; +/Δ, N=3, litter number=7; Δ/Δ, N=3, litter number=0) males are shown together. (D) Sperm counts from the epididymis of heterozygous (2.13 ± 0.17 cells) and homozygous (2.11 ± 0.09 cells) *CatSpert*-mutant males. Sperm numbers from *CatSpert-134del* (circles; +/Δ, N=5; Δ/Δ, N=5) and *128del* (triangles; +/Δ, N=3; Δ/Δ, N=3) are represented together. (E) Histology of *CatSpert* knockout testis. PAS-stained *CatSpert*^+/-^ (*left*) and *CatSpert*^-/-^ (*right*) testes sections are shown. (F) Computer Assisted Sperm Analysis (CASA) to measure sperm motility parameters from *CatSpert*^+/Δ^ (gray bars) and *CatSpert*^Δ/Δ^ (red bars) males. The motility parameters were measured from sperm cells before (0 min) and after (90 min) inducing capacitation. VCL, curvilinear velocity; ALH, amplitude of lateral head; VAP, average path velocity; VSL, straight line velocity; BCF, beat cross frequency. *\*p*<0.05 and *\*\*p*<0.01. N=3. (G) Flagella waveforms of sperm from *CatSpert-128del*^+/Δ^ (*top*) and *CatSpert-128del*^Δ/Δ^ (*bottom*) males. Sperm tail movements were recorded before (0 min, *left*) and after (90 min, *right*) capacitation. (H) Tail beating frequency of *CatSpert*^+/Δ^ (gray bars; 0 min, 11.26 ± 0.38 Hz; 90 min, 6.11 ± 0.72 Hz) and *CatSpert*^Δ/Δ^ (red bars; 0 min, 20.97 ± 2.28 Hz; 90 min, 26.13 ± 4.96 Hz) sperm. Circles indicate the tail beating frequency of individual sperm cells. N=45 from three animals for each group. *\*\*\*p*<0.001. (I) Immunoblotting of phosphorylated PKA substrates from *CatSpert*-mutant sperm cells. The extent of phosphorylation was compared between *CatSpert*^+/Δ^ and *CatSpert*^Δ/Δ^ sperm incubated under capacitating conditions (Cap) at the indicated times. Data is represented as mean ± SEM (C, D, F, and H).

**Figure S3.**
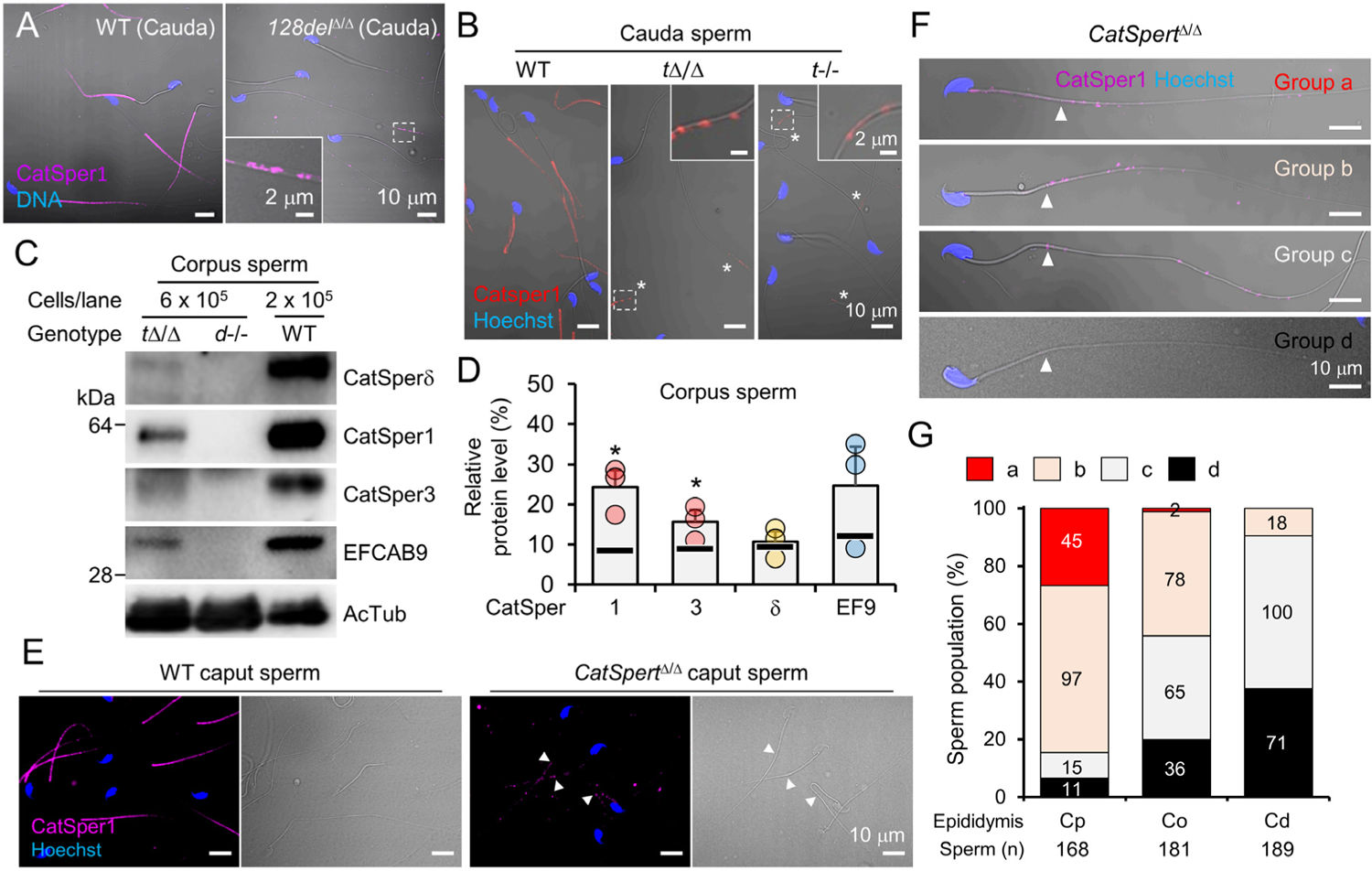
Diminished protein levels of CatSper subunits in epididymal *CatSpert*^D/D^ sperm, related to Figure 3. (A) Confocal images of the immunostained CatSper1 in the spermatozoa from cauda region of epididymis of *CatSpert-128del*^Δ/Δ^ males. Shown are CatSper1 (magenta) in WT (*left*) and *CatSpert-128del*^Δ/Δ^ (*right*) spermatozoa. (B) Confocal images of WT (*left*), *CatSpert*^Δ/Δ^ (*middle*), and *CatSpert*^-/-^ (*right*) cauda sperm immunostained with α-CatSper1. Asterisks (*) indicate immunolabelled CatSper1 in *CatSpert* mutant (*t*Δ/Δ) and knockout (*t*-/-) sperm. Magnified insets are shown (A and B). (C-D) CatSper subunit expression in *CatSpert*^Δ/Δ^ corpus sperm. (C) Immunoblotting of CatSper subunits in *CatSpert*^Δ/Δ^, *CatSperd*^-/-^ and WT corpus sperm. (D) Relative protein levels of CatSper subunits in corpus sperm from *CatSpert*^Δ/Δ^ compared to WT males. Each circle indicates relative protein levels of CatSper1 (24.2 ± 4.2%), CatSper3 (15.6 ± 3.0%), CatSperδ (10.6 ± 2.7%), and EFCAB9 (EF9, 24.7 ± 9.7%) in *CatSpert*^Δ/Δ^ corpus sperm compared to WT corpus sperm (N=3). Black lines in each column indicate the averages of the relative subunit levels in cauda sperm from *CatSpert*^Δ/Δ^ compared to WT males (see Figure 3B). Relative levels of CatSper1, 3, δ, and EFCAB9 in corpus and cauda *CatSpert*^Δ/Δ^ sperm were statistically compared. *\*p*<0.05. Data is represented as mean ± SEM. (E) Confocal images of WT and *CatSpert*^Δ/Δ^ caput sperm stained with α-CatSper1. (F-G) Different patterns of CatSper distribution in epididymal sperm from *CatSpert*^Δ/Δ^ males. (F) Four groups of the entire epididymal *CatSpert*^Δ/Δ^ sperm classified by distribution patterns of CatSper1 signals along the tail. Shown are images of immunostained CatSper1 in epididymal sperm from caput (groups a and b) and cauda (groups c and d). (G) Quantitation of sperm cells in each group (E) from caput (Cp), corpus (Co), and cauda (Cd) epididymis. Fluorescence and corresponding DIC images are merged. (A, B, and F). Arrowheads indicate annulus (E and F).

**Figure S4.**
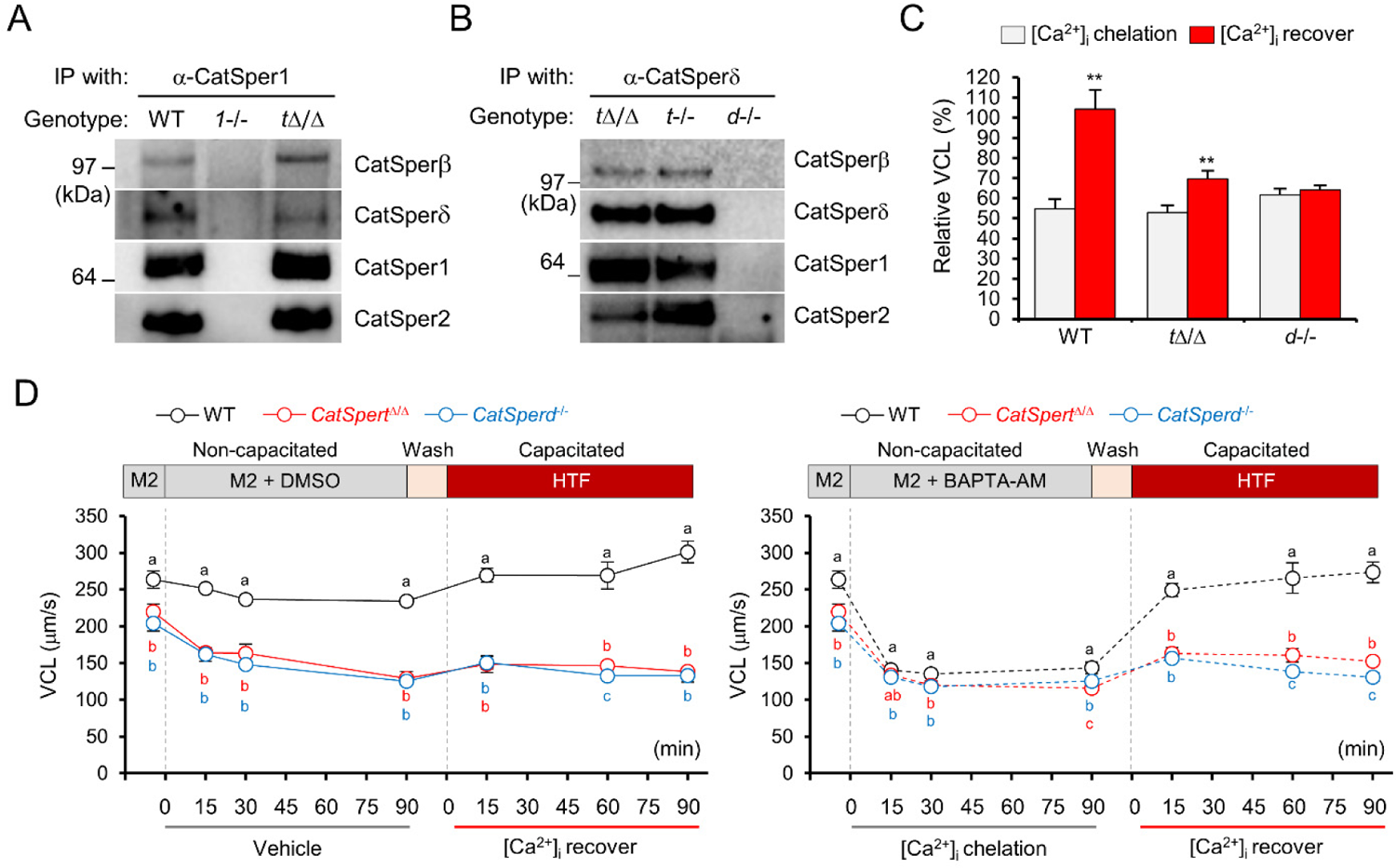
Marginal effects to CatSper channel complex assembly by CatSperτ malfunction, related to Figure 4. (A-B) Co-immunoprecipitation of CatSper complex with α-CatSper1 (A) and α-CatSperδ (B) in solubilized microsomes of *CatSpert*^Δ/Δ^ and *CatSpert*^-/-^ testis. WT and *CatSper1*-(A) or *CatSperd*-null (B) testes were used for positive and negative control, respectively. (C-D) Curvilinear velocity (VCL) changes by handling intracellular Ca^2+^ levels ([Ca^2+^]_i_) in WT, *CatSpert*^Δ/Δ^, and *CatSperd*^-/-^ spermatozoa. (C) VCL values of WT (black), *CatSpert*^Δ/Δ^ (red), and *CatSperd*^-/-^ (blue) sperm were measured during pre-incubation with vehicle (DMSO, *left*) or intracellular Ca^2+^ chelator (BAPTA-AM, *right*) in M2 and incubation in HTF medium. Different letters indicate significant differences in pairwise comparison between genotypes at each time point. (D) Relative VCL changes by [Ca^2+^]_i_ chelation (gray) and recovery (red). VCL values at the 90 minutes points from [Ca^2+^]_i_ chelation and recovery were normalized by the average VCL values before incubation with BAPTA-AM (before 0 min, M2) in each genotype. Statistical analyses were conducted by comparing relative VCL values after [Ca^2+^]_i_ chelation and recovery within each genotype. *\*\*p*<0.01. Data is represented as mean ± SEM (C and D). N=3.

**Figure S5.**
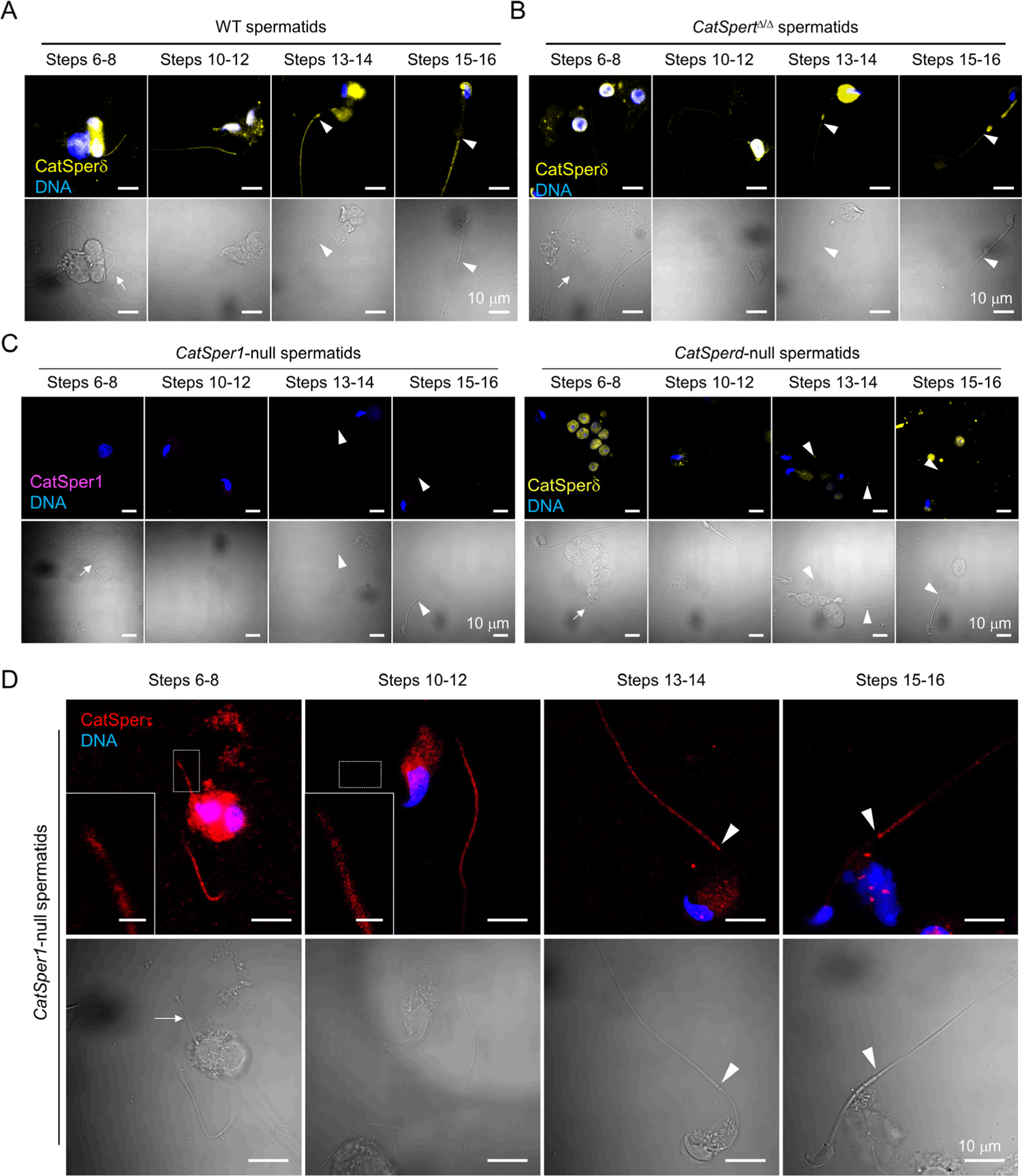
Targeting of CatSper proteins to the flagella and nanodomains in developing spermatids, related to Figure 5. (A-B) Confocal images of immunostained CatSperδ in WT (A) and *CatSpert*^Δ/Δ^ (B) spermatids. Fluorescence (*top*) and corresponding DIC (*bottom*) images and shown. (C) Confocal images of testicular spermatids from *CatSper1*^-/-^ (*left*) and *CatSperd*^-/-^ (*right*) males stained with α-CatSper1 (magenta) and α-CatSperδ (yellow), respectively. CatSperδ signal detected from the cytoplasm (steps 6-14), near annulus (steps 13-14), and mid-pieces (steps 15-16) of *CatSperd*^-/-^ spermatids are considered non-specific. Corresponding DIC images are shown (*bottom*). (D) Confocal images of CatSperτ-stained spermatids from *CatSper1*^-/-^ males. Corresponding DIC images of the immunostained *CatSper1*^-/-^ spermatids are shown (*bottom*). Insets are magnified (scale bar = 2 μm). Arrows and arrowheads indicate the elongating flagella of steps 6-8 spermatids and the annulus, respectively (A, B, C, and D). Hoechst were used for counter staining (A, B, C, and D).

**Figure S6.**
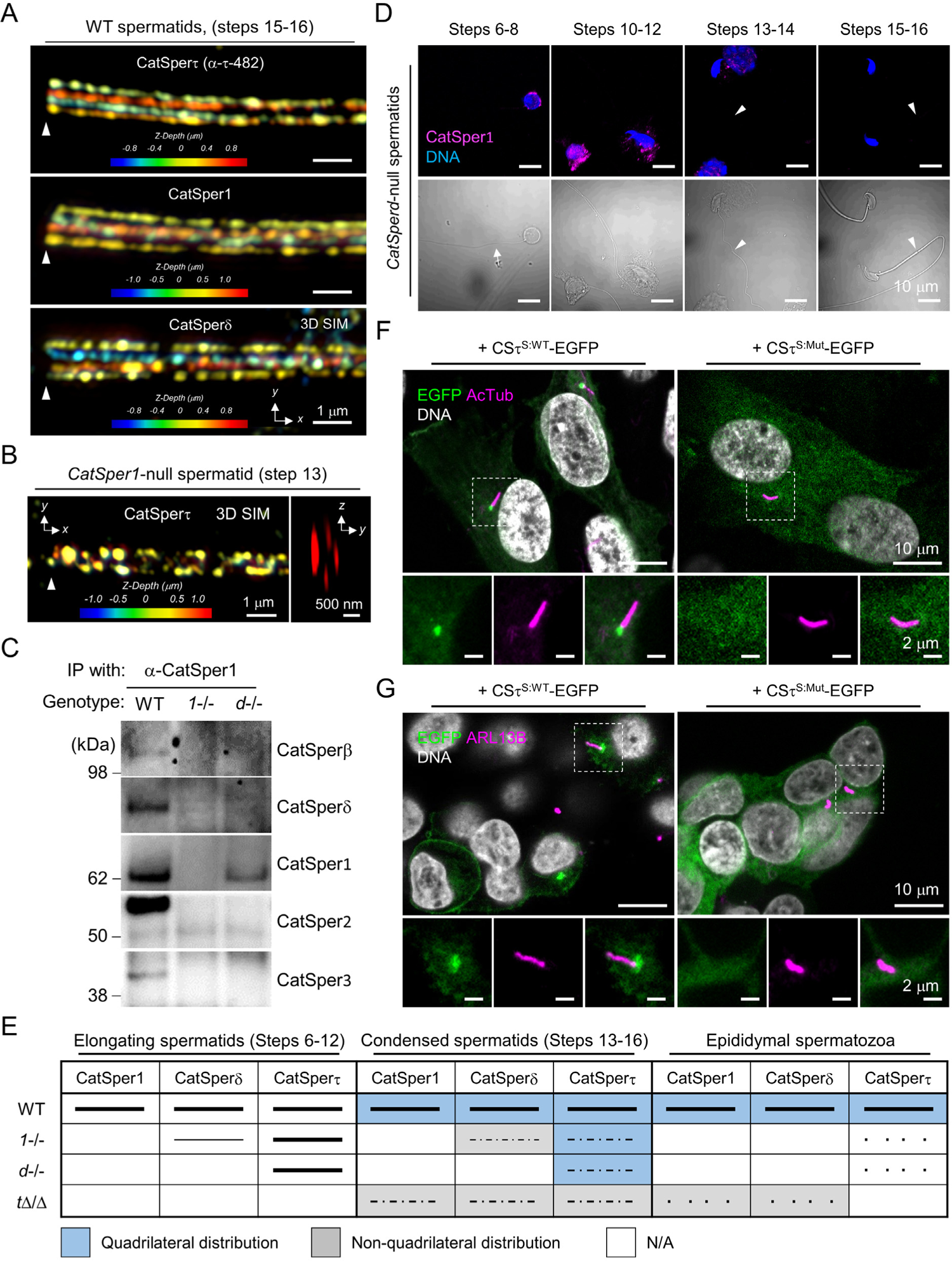
Capability of CatSperτ to traffic to the flagellum and organize quadrilinear nanodomains in developing spermatids, related to Figure 6 and 7. (A) 3D SIM images of CatSper subunits in WT spermatids at developmental steps 15-16. Shown are the images of immunostained CatSperτ (*top*), CatSper1 (*middle*), and CatSperδ (*bottom*) color-coded in *z*-depth from focal plane. (B) 3D SIM images of CatSperτ in *CatSper1*^-/-^ spermatids at steps 13-14. *x*-*y* projection is color-coded to represent *z*-depth from focal plane (*left*). *y*-*z* cross section is shown (*right*). (C) Co-immunoprecipitation of CatSper1 in *CatSperd*-null testis. None of the examined CatSper pore subunits (CatSper2 and 3) are detected from the CatSper1-immunocomplex in *CatSperd*^-/-^ testis (*d*-/-). WT and *CatSper1*-null (*1*-/-) testes were used for positive and negative control, respectively. (D) Confocal images of CatSper1 in developing *CatSperd*-null spermatids. CatSper1 is not detected in the elongating tails of *CatSperd*-null spermatids at all steps examined. Corresponding DIC images are shown (*right*). (E) Summary of the flagellar targeting and quadrilateral distribution of the CatSper proteins in male germ cells. Lines and colors in each cell depict arrangement and distribution of the CatSper proteins, respectively, in germ cells from each genotype. (F-G) Intracellular localization of the recombinant WT and the C2-domain truncated CatSperτ in ciliated hTERT-RPE (F) and 293T (G) cells. EGFP-tagged short forms of WT CatSperτ (CSτ^S:WT^-EGFP, *left*) and the predicted C2-truncated mutant CatSperτ (CSτ^S:Mut^-EGFP, *right*; Figure S1) were stably expressed in hTERT-RPE (F) and 293T (G) cells using lentiviral system. Immunostained acetylated tubulin (AcTub, F) and ARL13B (G) represent cilia (magenta). Enlarged insets are shown (*bottom*). An arrow (D) and arrowheads (A, B, and D) represent flagella in step 6-8 spermatids and annulus, respectively. Hoechst were used for counter staining (D, F, and G).

**Figure S7.**
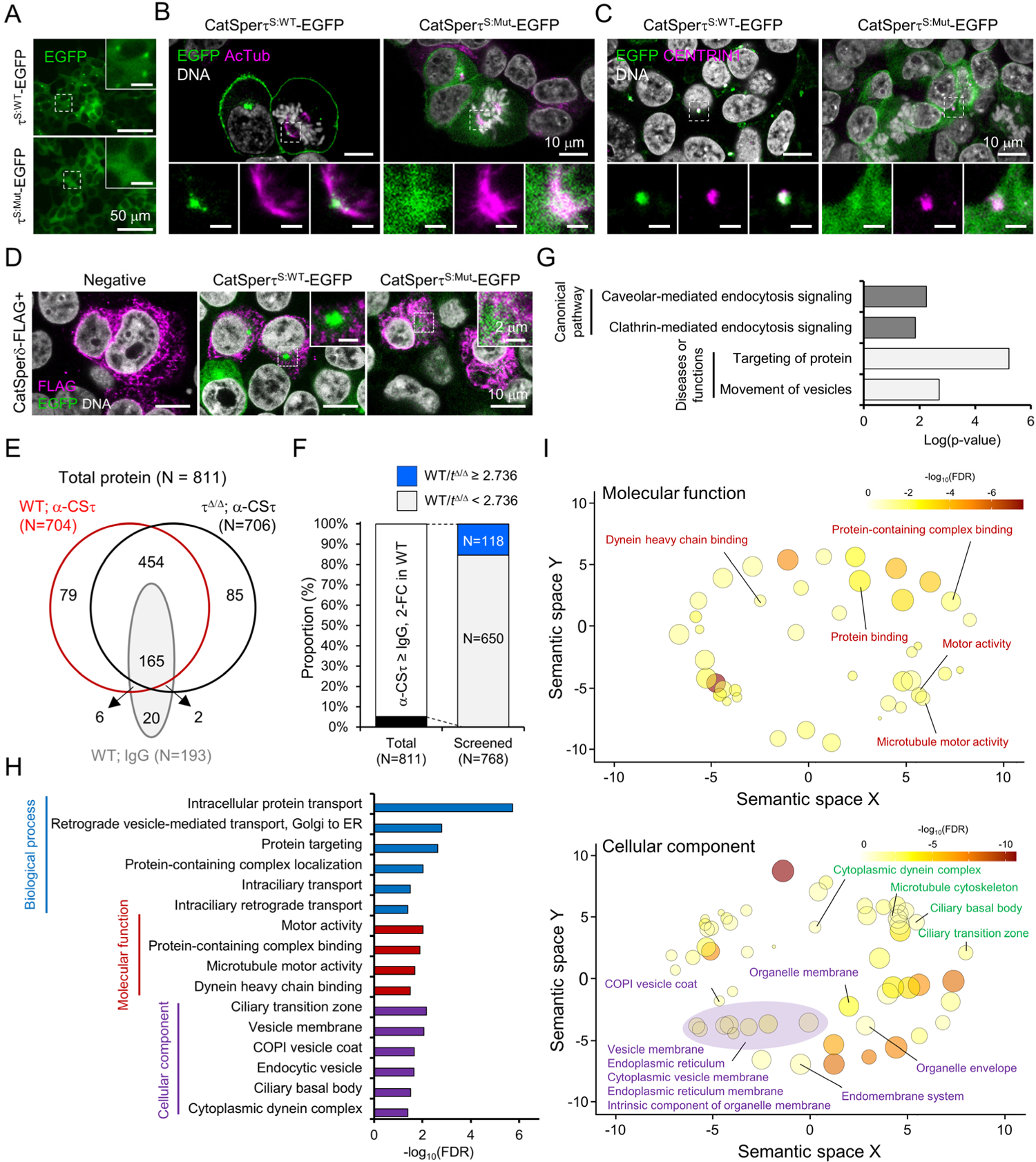
CatSpert is in complex with molecular motors and proteins in cytoplasmic vesicles, related to Figure 7. (A-C) Intracellular localization of recombinant CatSperτ in 293T cells. Epifluorescence (A) and confocal (B and C) images of short forms of WT (CatSperτ^S:WT-^ EGFP; A, *top*; B and C, *left*) and mutant (CatSperτ^S:Mut^-EGFP; A, *bottom*; B and C, *right*) CatSperτ expressed in 293T cells by lentiviral transduction (see also Figure 7A). EGFP is tagged at C-termini of the recombinant proteins. Immunostained acetylated tubulin (B, AcTub) and CENTRIN1 (C) show centrosomal localization of WT CatSperτ. Scale bars in magnified insets are 10 μm (A) or 2 μm (B and C). (D) Confocal images of immunostained CatSperδ in 293T cells expressing CatSperτ. FLAG-tagged CatSperδ (CatSperδ-FLAG) were transiently expressed in 293T cells that stably express WT (CatSperτ^S:WT^-EGFP) or mutant (CatSperτ^S:Mut^-EGFP) CatSperτ. 293T cells that are not transduced were used as negative control. Intracellular localization of CatSperδ is not changed by neither WT nor mutant CatSperτ in 293T cells. Magnified insets are shown (A, B, C, and D). Hoechst was used for counter staining (B, C, and D). (E-F) Number and proportion of the identified proteins from CatSperτ co-immunoprecipitation (coIP) mass spectrometry. (E) Venn diagram of total 811 proteins identified from three conditions of coIP mass spectrometry: coIP using α-CatSperτ-359 in WT (red, n=704) and *CatSpert*^Δ/Δ^ (black, n=706) testis, and coIP using rabbit normal IgG in WT testis (gray, N=193). (F) 768 proteins enriched more than 2-fold in the CatSperτ immunocomplex (white) compared to that from normal IgG (black) from WT testis (see also Figure 7E). Among them, 118 proteins including CatSperτ (blue) that showed more than 2.7-fold in WT over *CatSpert*^Δ/Δ^ were considered to CatSperτ-interactome, which were subjected to pathway analysis and functional annotation. (G-I) Enriched pathways (G) and gene ontologies (H and I) of the CatSperτ-interactome in testis. Canonical pathways (dark gray) and diseases or functions (light gray) from Ingenuity Pathway Analysis are shown (G). Representative biological process (blue), molecular function (red), and cellular component (purple) gene ontologies from functional annotation are listed (H). *X*-axis indicates negative logarithm of p value (G) or false discovery rate (FDR, H). (I) Molecular function (*top*) and cellular component (*bottom*) gene ontologies were functionally categorized. Enriched ontologies were represented as bubbles in scatter plot. Sizes and colors of the bubbles indicate percentage of proteins annotated with the ontologies and FDR, respectively.

**Video S1. Flagellar movement of tethered *CatSpert*^+/Δ^ and *CatSpert*^Δ/Δ^ sperm, related to** Figure 2 Uncapacitated (*top*, 0 min) and capacitated (*bottom*, 90 min) epidydimal sperm from *CatSpert*^+/Δ^ (*left*) and *CatSpert*^Δ/Δ^ (*right*) mice were tethered to imaging chamber and flagellar movement was recorded at 37 °C. Videos are played at 100 fps (1/2 speed).

Table S1. Identified proteins from CatSperτ interactome in testis by immunoprecipitation mass spectrometry, Related to Figure 7.

Table S2. Lists of the enriched pathways and gene ontologies of CatSperτ interacting proteins, Related to **Figure 7**.

